# A Multifaceted Framework To Assess tradeoffs in Interpretability, Explanatory and Predictive Performances Of Alternative Joint Species Distribution Models

**DOI:** 10.1101/2022.12.19.519605

**Authors:** Clément Violet, Aurélien Boyé, Mathieu Chevalier, Olivier Gauthier, Jacques Grall, Martin P. Marzloff

**Affiliations:** IFREMER, Centre de Bretagne, DYNECO LEBCO, Plouzané, France; Laboratoire des Sciences de l’Environnement Marin (LEMAR) UMR 6539 CNRS UBO IRD IFREMER, Institut Universitaire Européen de la Mer, Université de Bretagne Occidentale, Plouzané, France; Observatoire des Sciences de l’Univers, UMS 3113, Institut Universitaire Européen de la Mer, Plouzané, France

**Keywords:** Community assembly, Joint Species Distribution Model, Explanatory power, Predictive power, Interpretability, Model Performances

## Abstract

*Joint Species Distribution Models* (*jSDM*) are increasingly used to explain and predict biodiversity patterns. By accounting for species co-occurrence patterns and potentially including species-specific information, *jSDM*s capture the processes that shape ecological communities. Yet, factors like missing covariates or omitting ecologically-important species may alter the interpretability and effectiveness of *jSDM*s. Additionally, while the specific formulation of a *jSDM* directly affects its performances, the effects of choices related to model structure, such as inclusion, or not of phylogeny or trait information, are not well-explored.

Here, we developed a multifaceted framework to comprehensively assess performances of alternative *jSDM* formulations at both species and community levels. We applied this framework to four alternative models fitted on presence/absence and abundance data of a polychaete assemblage sampled in two coastal habitats over 500 km and 8 years. Relative to a benchmark *jSDM* only capturing the effects of abiotic predictors and residual co-occurrence patterns, we explored the performance of alternative formulations that also included species phylogeny, traits, or some additional 179 non-target species, which were sampled alongside the species of interest. For both presence/absence and abundance data, explanatory power was good for all models but their interpretability and predictive power varied. Relative to the benchmark model, predictive errors on species abundances decreased by 95% or 53%, when including non-target species, or phylogeny, respectively. These differences across models relate to changes in both species-environment relationships and residual co-occurrence patterns. While considering trait data did not improve explanatory or predictive power, it facilitated interpretation of trait-mediated species response to environmental gradients.

This study demonstrates trade-offs in *jSDM* formulation for explaining or predicting species data, highlighting the importance of using a comprehensive framework to compare models.

Furthermore, our study provides some guidance for model selection tailored to specific objectives and available data.

## Introduction

Community ecology aims at describing, explaining, and predicting changes in communities (Tredennick et al. 2021). Understanding the processes that determine species distribution is a prerequisite to characterise and predict community structure and associated ecological dynamics, which is critical to inform effective management or restoration measures in a rapidly changing world [Dietze et al. (2018); Brudvig and Catano (2022). *Joint Species Distribution Models* (*jSDM*) are particularly well-suited tools to address these challenges, whether to characterise the processes that shape observed communities (Warton et al. 2015; Ovaskainen, Tikhonov, Dunson, et al. 2017), or to predict future changes in species assemblages (Norberg et al. 2019; Pollock, O’Connor, et al. 2020).

*jSDM*s are multivariate (i.e. multi-species) extensions of *Species Distribution Models* (*SDM*s), which have been broadly applied over the past decades - across both terrestrial and marine realms - to understand and predict species occurrences (Elith et al. 2006; Norberg et al. 2019) and/or abundances (Howard et al. 2014; Waldock et al. 2022) using a set of covariates (e.g. climatic variables) as predictors. Relative to *SDM*s, *jSDM*s explanatory power can benefit from accounting for species-specific assembly rules (Ovaskainen, Tikhonov, Norberg, et al. 2017). In particular, relative to single-species *SDM*s that only consider the abiotic niche of species (i.e. the Grinellian niche), *jSDM* can theoretically also account for interspecific interactions (i.e. the Eltonian niche).

In *jSDM*s, the variability in community composition not explained by covariates is captured by a residual covariance matrix representing species co-occurrence patterns potentially capturing biotic interactions (Ovaskainen, Tikhonov, Norberg, et al. 2017). This feature is, at first glance, highly attractive to ecologists because it provides a way to disentangle the relative influence of abiotic and biotic processes on biodiversity patterns (Godsoe, Franklin, and Blanchet 2017) while also improving model’s predictive power (Giannini et al. 2013; Staniczenko et al. 2017). However, in practice, inferring and interpreting residual co-occurrence patterns using *jSDM*s remains challenging for several reasons (Blanchet, Cazelles, and Gravel 2020; Holt 2020).

First, while *jSDM*s have been applied to a large number of species presence/absence datasets (Norberg et al. 2019; Wilkinson et al. 2019; Wilkinson et al. 2021), simulation studies showed that inferred co-occurrence networks do not necessarily provide evidence for species interactions (Dormann et al. 2018; Blanchet, Cazelles, and Gravel 2020) but only capture spatial and temporal associations between species (Keil et al. 2021). Some authors speculated that *jSDM*s applied to abundance data - instead of presence/absence data - could provide a better proxy for biotic interactions (Blanchet, Cazelles, and Gravel 2020; Momal, Robin, and Ambroise 2020). Accordingly, *jSDM*s have increasingly been applied to abundance data (Hui 2016; Ovaskainen, Tikhonov, Norberg, et al. 2017; Chiquet, Mariadassou, and Robin 2021). While challenges related to modelling abundance data was recently explored in the context of single species distribution modelling (*SDMs*; Waldock et al. (2022)], the predictive and the explanatory power of *jSDM*s fitted to abundance data remains relatively untested compared to presence/absence data (Norberg et al. 2019; Wilkinson et al. 2021). Second, regardless of the type of data considered (i.e. presence/absence or abundance), several factors may limit or affect the interpretability and predictive ability of *jSDM*. For instance, co-occurrence patterns estimated in *jSDM* are affected by unaccounted environmental variables im-plying that *jSDM*s cannot fully separate the environmental and the biotic niche of species (Blanchet, Cazelles, and Gravel 2020; Poggiato et al. 2021). Beyond missing environmental predictors, accounting for extra species that can influence the target community (e.g. competitors) is key to improve *jSDM*s’ inference and predictions. However, because many ecological studies only focus on particular taxonomic groups (Pollock, Tingley, et al. 2014; Häkkilä et al. 2018) and disregard non-target taxa, co-occurrence patterns estimated from *jSDM*s are almost always skewed by missing ecological actors (Momal, Robin, and Ambroise 2021). How this bias affects the predictive ability of *jSDM*s remains untested.

Finally, similar to *SDM*s, *jSDM*s can theoretically be extended to include additional sources of information about modelled species (Niku et al. 2019; Ovaskainen, Tikhonov, Norberg, et al. 2017). For instance, accounting for phylogenetic relationships between species (Ives and Helmus 2011) or for the link between functional traits and environmental responses (Pollock, Morris, and Vesk 2012) can improve both the explanatory and the predictive powers of *SDM*s (Morales-Castilla et al. 2017; Vesk et al. 2021). These findings support the hypothesis that similar species, in terms of traits and/or recent evolutionary history, usually share similar environmental preferences. While inclusion of species-specific information in *jSDM*s should yield similar benefits (Ovaskainen, Tikhonov, Dunson, et al. 2017), the relative influence of additional sources of information on their interpretability and predictive power remains untested (Norberg et al. 2019; Wilkinson et al. 2019; Abrego, Bässler, et al. 2022).

Overall, many practical questions remain concerning the application of *jSDM*s to ecological community monitoring data, in particular related to the inclusion of additional sources of information within the models. While some comparative assessments of *jSDM*s performance exists (e.g. Norberg et al. (2019); Wilkinson et al. (2021)), including some comparison of the benefit of trait and phylogenetic data in some phyla (e.g. Abrego, Bässler, et al. (2022)), there has been no formal assessment of the relative importance of species-specific information (trait and/or phylogeny) compared to the role of missing species. Furthermore, comparative assessments have rarely been performed on both presence/absence and abundance data. To a few exceptions (Waldock et al. 2022), most assessments were made considering presence/absence datasets (Norberg et al. 2019; Wilkinson et al. 2019) and mostly focused on predictive power (Norberg et al. 2019; Wilkinson et al. 2019), hence disregarding the interpretability/explanatory aspects of the models (Tredennick et al. 2021). Yet, *jSDM*s are increasingly used for explanatory purposes (Abrego, Dunson, et al. 2017; Jacobi and Siqueira 2023).

Hence, there is a mismatch between the current understanding of *jSDM*s performances and their implementation by ecologists. In practice, most *jSDM* applications consider a single model structure and do not explore the effects of including additional sources of information to answer their specific research question. Perhaps this shortcoming relates to the high-dimensionality of *jSDM*s which makes their comparison challenging without a proper multi-faceted assessment framework.

In this study, we developed such a framework to evaluate the extent to which alternative parameterisation of *jSDM* can lead to a better interpretability or predictability at species and community levels. To illustrate its usefulness, we applied this general framework to a case study presenting typical features of community ecology datasets. Specifically, by comparing predictions obtained from a *Benchmark* model excluding additional sources of information (i.e. a classical *jSDM*), we tested the effect of (1) including phylogeny alone and in combination with trait data, (2) incorporating monitoring information related to non-target species and (3) considering abundance instead of presence/absence data. We hypothesised that these additional data (i.e. phylogeny, traits and non-target species) should improve *jSDM* predictive and explanatory powers at species and community levels, but did not assume a priori that a given modelling strategy would lead to greater improvements in model performances at either level of biological organisation.

## Materials & Methods

We used the *HMSC* (*Hierarchical Modeling of Species Communities*) framework applied to a long-term monitoring dataset. The following subsections sequentially describe our workflow (as illustrated in fig. 1): (1) the *HMSC* framework and (2) the case study specifics Subsections 3-5 details how to implement our multi-facetted framework in a stepwise manner, namely in terms of: (3) data splitting between train and test datasets to assess the explanatory and predictive powers of models, respectively, (4) definition of the suite of relevant alternative models and, (5) a multi-faceted framework developed to assess tradeoffs in *jSDM*s’ performances in relation to different study purposes.

**Figure 1.**
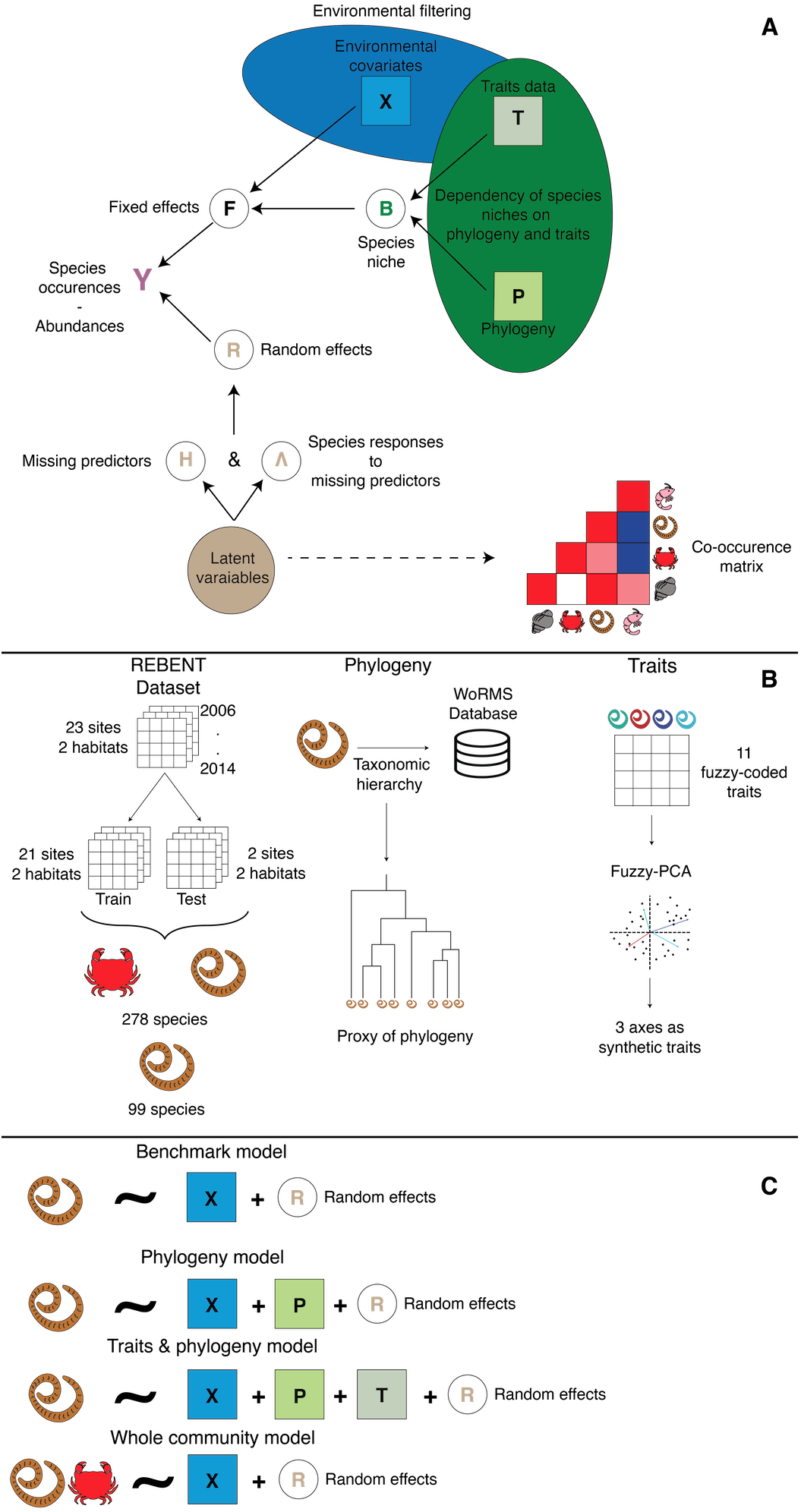
Study workflow. A. Structure of the *Hierarchical Model of Species Community* (*HMSC*), incorporating environmental variables, phylogeny, and species-specific traits. B. Data pre-processing: partitioning community data into train and test datasets, estimating phylogenetic distance (using taxonomic classification), and dimension reduction of the species-trait matrix using a fuzzy-PCA. C. Summary of the four alternative model structures fitted to presence/absence and abundance data: the *Benchmark*, *Phylogeny*, and *Traits & Phylogeny* models only consider a target assemblage of 99 polychaetes, while the Whole Community model includes additional species (179) potentially interacting with the target assemblage. Random effects for sampling years, sites, and habitats were included in all models.

### Hierarchical Modelling of Species Community (HMSC)

*HMSC* is a multivariate hierarchical generalised linear mixed model using Bayesian inference (Ovaskainen and Abrego 2020). It has two parts: one for fixed effects and another for random effects. The fixed part models the species’ realised niche, where each dimension of the niche is a covariate (e.g. temperature; fig. 1). Including trait data can improve species niche estimates by accounting for trait-environment relationships, where species with similar traits are expected to respond similarly along environmental gradients (Ovaskainen, Tikhonov, Dunson, et al. 2017). Including phylogenetic data can help capture residual ecological information not included in the available trait data, as phylogenetically-close species tend to share similar traits and niches (Wiens et al. 2010). Alongside traits, phylogeny can improve niche estimates for rare species by borrowing information from similar species (Ovaskainen, Tikhonov, Dunson, et al. 2017; Ovaskainen, Tikhonov, Norberg, et al. 2017; Ovaskainen and Abrego 2020). The random part of *HMSC* relies on latent variables, i.e. covariates that capture residual variance, including missing environmental features or biotic interactions (Ovaskainen, Tikhonov, Dunson, et al. 2017; Ovaskainen, Tikhonov, Norberg, et al. 2017; Ovaskainen and Abrego 2020). The *H* matrix (site loadings) estimates missing covariate values, while the Λ matrix (species loadings) represents species’ responses to these missing covariates (fig. 1).

### Case Study

#### Faunistic data

The REBENT (rebent.ifremer.fr) is an ongoing monitoring program of benthic macrofauna performed across multiple stations along Brittany’s coastlines (Western France). Here, we used data from Boyé, Thiébaut, et al. (2019), covering 23 sites and two intertidal soft-bottom habitats: bare sands and seagrass meadows (Fig. S1) where infaunal communities were monitored yearly using the same protocol between 2006 and 2014. A detailed description of the sampling methodology is provided in (Boyé, Legendre, et al. 2017; Boyé, Thiébaut, et al. 2019). At each site, three faunal samples (0.03 m^2^ cores) were taken at each of the three fixed sampling points distributed 200 m apart. These samples were then pooled together to estimate abundances at the site level. For each sampling event, individuals were identified to the lowest taxonomic level possible (mostly species level; for simplicity we hereafter refer to “species”).

Overall, across a total of 375 sampling units (i.e. unique combination of years, sites and habitats), 152,583 individuals belonging to 519 species were collected and identified. To avoid convergence issues and poor model inference, we kept only species occurring at least four times across the 180 samples used as train set (see below), resulting in the removal of 241 species. The remaining 278 species included 99 polychaete species (the target assemblage) and 179 non-target species of bivalves, molluscs, and amphipoda, which may predate or compete with polychaetes (Grall et al. 2006; Jankowska et al. 2019). We chose to focus on polychaetes as this taxonomic group exhibits diverse lifestyles (Jumars, Dorgan, and Lindsay 2015), can be used to monitor the health of benthic habitats (Giangrande, Licciano, and Musco 2005), and because trait data and ecological information were available from previous studies (Boyé, Thiébaut, et al. 2019).

### Traits and phylogeny data

Traits data were retrieved from (Boyé, Thiébaut, et al. 2019) for the 99 polychaete species present in the train set (see below). The 11 fuzzy-coded traits available (see Boyé, Thiébaut, et al. (2019) for details) were synthesized using a fuzzy-PCA, with the *fpca* function from the *ade4* R package (Thioulouse et al. 2018). The first three axes, which account for 59% of the total variance of the trait matrix, were included in the model as synthetic traits data (Fig. 1). The first axis distinguishes mobile predatory species from sessile microphages whereas the second axis differentiates semelparous species from iteroparous species. The third axis separates burrowers from tube-dwellers (Fig. S2). Phylogeny was not available; hence we followed common practices (Ovaskainen & Abrego 2020) and retrieved the taxonomic classification of these 99 polychaetes through the WoRMS database (www.marinespecies.org; January 2020) and used this information as a proxy for phylogenetic relationships (fig. 1; Ovaskainen and Abrego (2020)). Phylogenetic distances were then estimated using the *ape* R package (Paradis and Schliep 2019).

### Environmental data

Following Boyé, Thiébaut, et al. (2019) (see Appendix A for details about data sources), we selected seven environmental variables to characterise the ecological niche of each species. These seven variables quantify different components of coastal environmental variability including hydrology (sea water temperature, salinity and current velocity), sedimentology (mud and organic matter content), substrate heterogeneity (Trask index) and local wave exposure (fetch). For each of these seven variables, the first- and second-degree polynomials were computed to account for non-linear responses.

### Stepwise Implementation of a multi-facetted assessment framework

#### Comparison of alternative model structures

The first model (denoted *Bench*) only considers data for the 99 target polychaete species and the seven environmental covariates (fig. 1). The second model (denoted *Ph*) adds phylogenetic data to the *Bench* model (fig. 1). The third model (denoted *TrPh*) adds traits data to the *Ph* model. The fourth model (denoted *WhC*) has the same structure as the *Bench* model but includes data on the whole community (278 species, including 179 additional non-target species; fig. 1). *WhC* excludes traits (unavailable for the non-target taxa) and phylogenetic data for faster computation. Each model was fitted twice, either with presence/absence or abundance data, using probit and lognormal Poisson distributions respectively. All models include the same random effects (fig. 1): temporal (years), spatial (sites), and habitats (bare vs seagrass).

### Model fitting

We estimated model parameters by running 5 chains using MCMC sampling over 375,000 iterations. The first 125,000 iterations were discarded as burn-in while the remaining 250,000 iterations were thinned every 250 iterations yielding 1,000 posterior samples per chain for an overall total of 5,000 posterior samples for each parameter. We assessed convergence for each model parameter using both potential scale reduction factor (Gelman and Rubin 1992) and effective sample size (see supplementary materials, Appendix B). All models were fitted using the DATARMOR supercomputing facility.

### Assessing model performance and interpretability

For independent assessment of models’ predictive performance, the dataset was split into a train and a test set, instead of using a strict cross-validation procedure that would have considerably increased the computational burden (see also Norberg et al. (2019)). The train dataset consisted of 180 sampling units (21 sites; one or two habitats and six to nine years per site; Fig. S1). The test set comprised 35 sampling units covering a 9-year period at two specific sites comprising both seagrass and bare sand habitats. These sites were chosen as representative of both regional macrofaunal species diversity (all the species observed in the test set are also observed in the train set) and mean environmental conditions (which limits model extrapolation outside of the trained parameter space; Fig. S3-S4; Boyé, Legendre, et al. (2017); Boyé, Gauthier, et al. (2022); Toumi et al. (2023)). Another argument was the presence of monitoring stations in both habitats in these two sites, which was necessary to assess potential habitat-related biases in model predictions.

To assess *jSDM*’s performance, we used a set of complementary metrics to evaluate both their explanatory and predictive abilities on the train and test dataset, respectively (Table 1). All metrics were computed only for the target assemblage (i.e. polychaetes) - even for the *WhC* model that includes a total of 278 species - to ensure a proper comparison between models. AUC and RMSE were used to assess the overall and species-level performance for presence/absence and abundance models, respectively. Relationships between observed and predicted average species abundances across all sites were also visualised for abundance models.

**Table 1.**
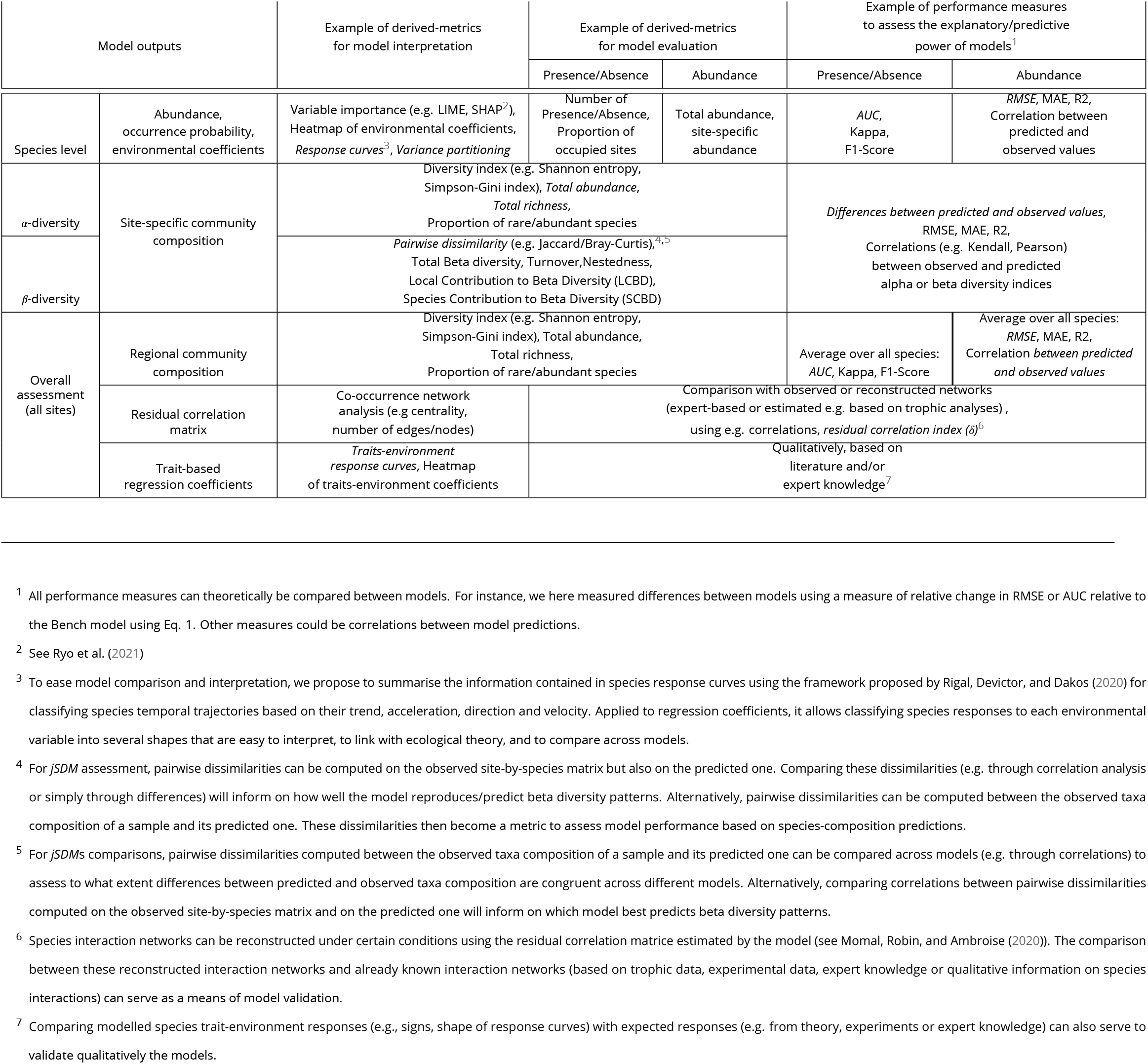
Multi-assessment framework providing a list of useful metrics to assess, interpret or compare jSDMs across different ecological facets (rows) at the species, community or overall level. Italicized metrics are used in this study.

Along with raw AUC and RMSE values, we also visualised and quantified changes relative to the *Bench* model for the *Ph*, *TrPh* and *WhC* models. For abundance models, we computed the overall relative change in mean RMSE across species as:

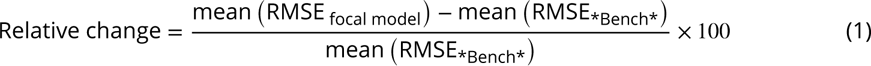

AUC and RMSE only partially capture model accuracy at the community scale (Table 1). To explore this aspect, we focused on differences between predicted and observed assemblage richness and total abundances (for abundance models). We also compared observed and predicted Sørensen (for presence/absence) and Bray-Curtis (abundance) pairwise-dissimilarity matrices to explore how well β-diversity patterns were reproduced by the models. For these three metrics, we computed relative change for both the train and test datasets between mean predicted and mean observed values as follows:

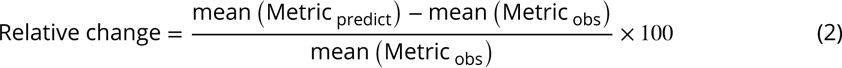

where “Metric” is a community-based measure (e.g. species richness, total abundance, dissimilarity matrices) estimated from model predictions or observations at the sample level (i.e. unique combination of site, habitat and year or pairs of samples for dissimilarity). To evaluate model interpretability, we calculated the amount of explained variance per species and the proportion that can be attributed to environmental covariates (fixed effects) and random effects. We compared the overall relative change in the proportion of variance explained by the covariates and by the random effects for the *Ph*, *TrPh* and *WhC* relative to the *Bench* model (by comparing mean values across species similarly to eq. 1). We also propose a novel way of exploring species-environment relationships (Table 1) by classifying the response curves estimated from the different models based on their shapes, considering both their direction (decline, null, or increase) and their acceleration (decelerated, constant, or accelerated; Rigal, Devictor, and Dakos (2020)). Finally, we compared the residual co-occurrence patterns associated with each random effect of the *Bench* model with those of the best performing model (*WhC*). We quantified differences in magnitude and sign of residual species-species correlations using the following index:

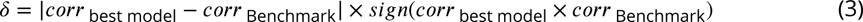

## Results

Both MCMC convergence and effective sample size of the different *jSDM*s were satisfactory (see Appendix D).

### Model Fit & Predictive power

#### Species level

Presence/absence models showed excellent explanatory power with mean AUCs above 0.9 on the train dataset, but lower predictive power with mean AUCs around 0.66 on the test set (Fig. S17). Both explanatory (mean AUC between 0.92 and 0.93) and predictive (mean AUC between 0.64 and 0.66) powers were similar across models (fig. 2, Fig. S17). Within the target species assemblage, predictions improved for 41 species and worsened for 36 species, out of the 99 target species (no changes for the remaining 22 species) for the *WhC* model relative to the *Bench* model leading to an overall improvement of 1.5% in AUC. In comparison for the *Ph* or the *TrPh* models, predictions only improved for 26 and 28 species, and worsened for 49 and 46 species respectively, leading to an overall decrease in AUC of 0.3% and 0.1% respectively.

**Figure 2.**
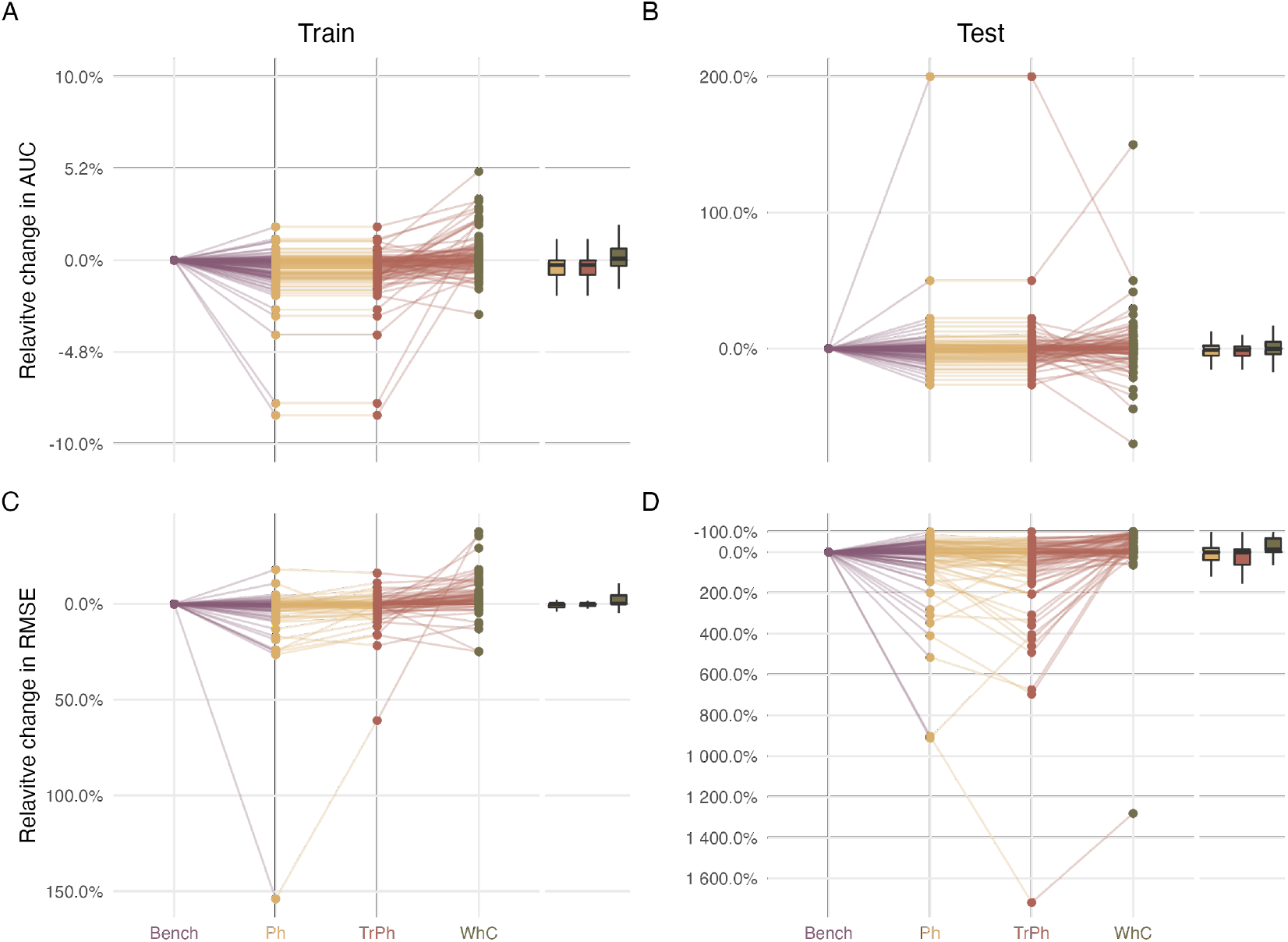
Change in explanatory and predictive power of three model structures (yellow for *Phylogeny* (Ph), red for *Traits and phylogeny* (TrPh), and green for *Whole community* (WhC) models) relative to the *Benchmark model* (Bench; purple). Changes are expressed as percentages relative to the benchmark fitted on presence/absence (top row) or abundance (bottom row) data. Points above the zero line indicate performance gain.

Abundance models also showed a satisfactory explanatory power with a mean RMSE close to nine for all models, given a mean abundance in the train dataset of 307.3 ± 583.6 (mean ± sd). Overall, all models underpredicted species abundances (Fig. S18-19). While explanatory power was similar across models, larger variations were observed for predictive power. The *Bench* model had a mean RMSE of 129.4 (for a mean abundance in the test dataset of 700.57 ± 818.66; Fig S17). The *Ph* model performed better with a mean RMSE of 58.4 (54.9% decrease in RMSE compared to the *Bench*; Fig. S17) whereas the *TrPh* model did worse with a mean RMSE of 139.2 (9.9% increase in RMSE compared to the *Bench*; Fig. S17). The best model was the *WhC* with a mean RMSE of 6.4 (95.0% decrease in RMSE compared to the *Bench*, Fig. S17). Out of the 99 target species, the *WhC* model predictions improved for 59 species (average decrease in RMSE = 52%) but declined for 15 species (average increase in RMSE of 105%) relative to the *Bench*, leading to an overall decrease in RMSE of 15.5%. Conversely, performance gain for the *Ph* and *TrPh* models were poor relative to the *Bench*, as predictions improved for 39 and 30 species (average decrease in RMSE of 33% and 32%, respectively), but declined for 39 and 45 species (average increase in RMSE of 137% and 179%, respectively), leading to an overall increase in RMSE of 41. 0% and 71.5% respectively.

We further investigated the gain in predictive power observed for the *WhC* model fitted to abundance data by examining the relationships between changes in predictive power and the occurrence or abundance of the species. Overall, we found no clear patterns between RMSE change and average species abundance (Kendall’s τ = 0.13, p-value = 0.07; Fig. S20) nor the proportion of presence (Kendall’s τ = 0.11, p-value = 0.14; Fig S21).

### Community level

In terms of alpha diversity, the *Bench*, the *Ph* and *TrPh* models fitted on abundance data all underpredicted the species richness of the train set by 4 species on average (−29.2% compared to observed data; Fig. 3). In contrast, the *WhC* model overpredicted species richness by 11 species on average (+80% compared to observed data). Similar results were found on the test dataset with the *Bench*, *Ph* and *TrPh* models underpredicting richness by 5 species on average (−24.9 %) whereas the *WhC* model overpredicted richness by 7 species (+35.8% compared with observed data). Similar results were found for models fitted on presence/absence data (Fig. S22).

**Figure 3.**
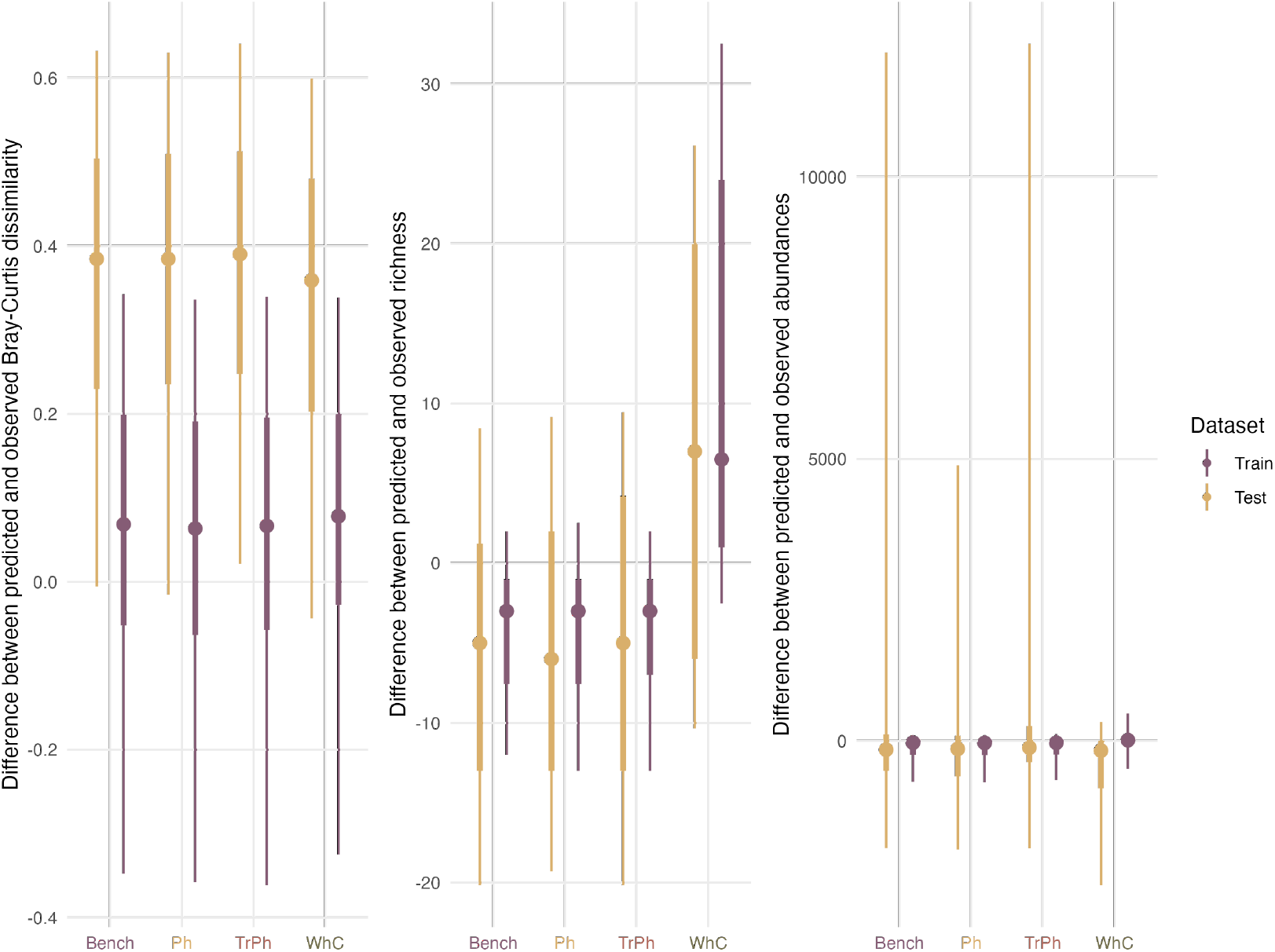
Comparison of model performances with regards to their ability to predict community structures when fitted with abundance data for the train (purple) and test (yellow) dataset. Left: differences in pairwise dissimilarities estimated on the observed and the predicted assemblages. Centre: differences in species richness between observed and predicted assemblages. Right: differences in total abundance between observed and predicted assemblages.

All models underpredicted total abundance relative to the train dataset on average (Fig. 3). The *Bench* model underpredicted total abundance by 153 individuals (−49.8% compared to observed data), while the *Ph* and *TrPh* models underpredicted by 159 and 155 individuals (−51.7% and - 50.4%), respectively. The *WhC* model only underpredicted total abundance by 22 individuals (−7.12% compared to observed data). Relative to the test dataset, the *Bench*, the *Ph* and the *TrPh* models overpredicted total abundance by 1642 (+234% compared to observations), 465 (+66.3%), and 1969 (+281%) individuals on average, respectively. In contrast, the *WhC* model underpredicted total abundance by 404 individuals on average (−57.6%).

Beta diversity patterns were overall well captured by all models fitted on abundance or presence/absence data regarding the train dataset (fig. 3). Observed dissimilarities were slightly over-predicted by all abundance models: by 0.057 for the *Bench* (+7.3% compared with observed data), 0.050 for the *Ph* (+6.4%), 0.054 for the *TrPh* (+6.9%) and 0.070 for the *WhC* models (+8.9%). Differences for presence/absence models were of similar order but all models underpredicted pairwise dissimilarities between samples on average (Fig. S22). On the test dataset, beta diversity patterns were poorly captured by the models fitted on abundance data. The *Bench* model overpredicted pairwise dissimilarities by 0.364 on average (+67.1% compared with observed data), the *Ph* model by 0.365 (+67.4%), the *TrPh* model by 0.375 (+69.1%) and the *WhC* model by 0.338 (+62.4%). Similar results were observed for presence/absence models with slightly smaller overpredictions (Fig. S22).

### Variance partitioning

The amount of variance explained across the 99 polychaetes varied between 21% and 23% for models fitted with presence/absence data and between 18% and 30% for abundance-based models (Fig. S23). For all models, environmental variables, rather than random effects, accounted for most (> 68% ± 18%; mean ± s.d.) of the explained variance (Fig. S24) except for the *WhC* model where a larger part of the variance was explained by random effects (Fig. S24). Compared to the *Bench* model fitted with abundance data, the relative change in the part of variance explained by random effects decreased by 17.00% for the *Ph* model, 10.90% for the *TrPh* model and increased by 224% for the *WhC* model (fig. 4). Similar results though with smaller relative changes were obtained for presence/absence models (Fig. 4; Fig. S23-24).

**Figure 4.**
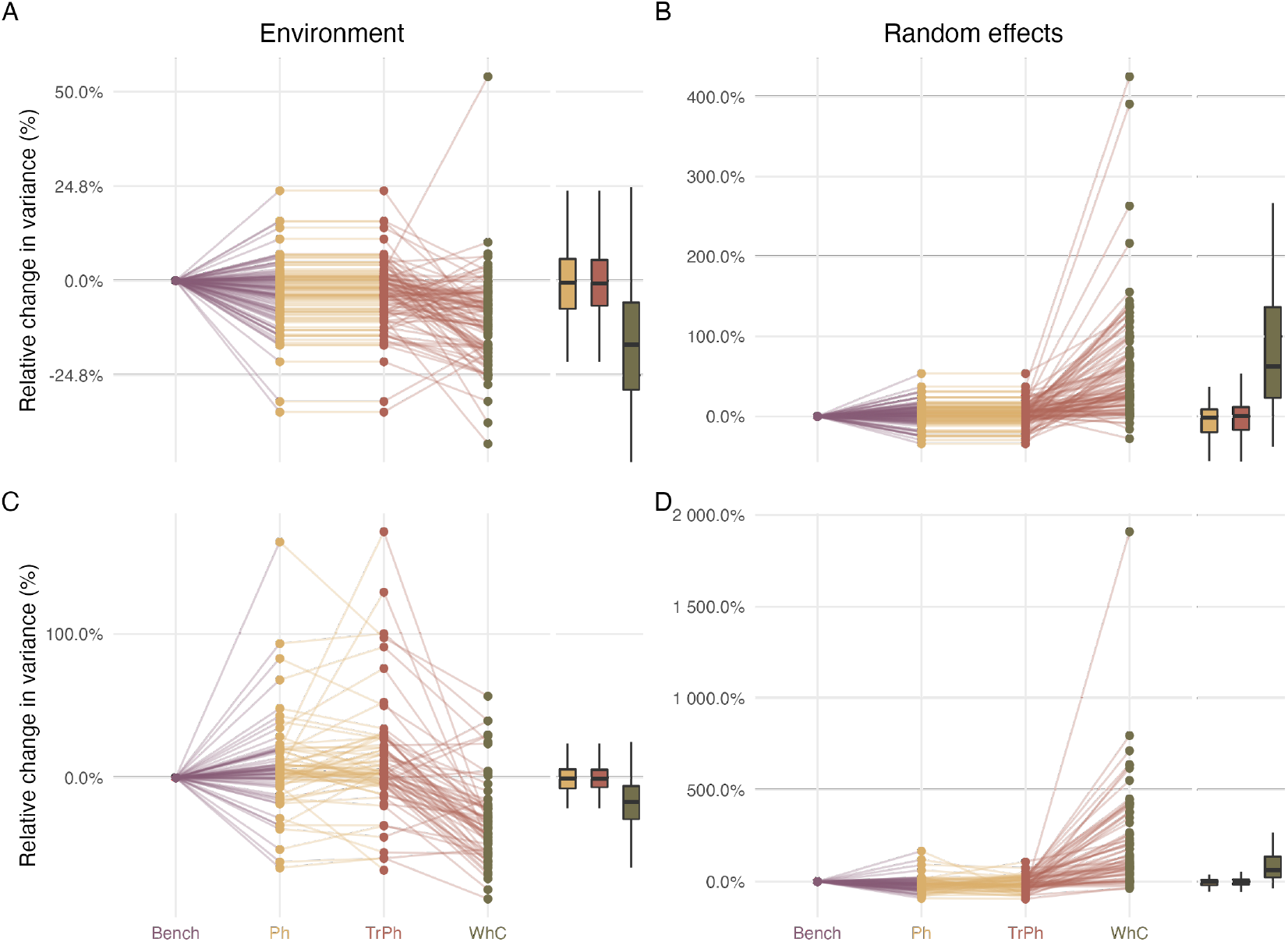
Change in explained variance related to environmental predictors (left column) and random effects (right column) for three alternative model structures (yellow for Phylogeny (*Ph*), red for Traits and phylogeny (*TrPh*), and green for Whole community (*WhC*) models) relative to the Benchmark model (*Bench*; purple). Percentage changes were computed relative to the *Bench* model fitted with presence/absence (top panels) or abundance (bottom panels) data. Positive values indicate an increase in the proportion of variance explained by the focal model compared to the *Bench* model. See Figure S23 and S24 for the raw percentages, expressed as percentages of explained variance or total amount of variance, respectively.

### Species niche estimated

For abundance models, the large majority (>60%) of response curves were flat indicating a lack of meaningful species-environment relationships (fig. 5). This proportion reached 83% for the *WhC* model. The prevalence of flat relationships did not appear to be related to convergence issues (Fig. S15-16) or to be driven by a specific covariate (Fig. S25). Convex or concave response curves were rare in abundance models. Significant relationships primarily included constant or accelerated declines, representing approximately 10% and 15% of response curves in the *Bench*, *TrPh*, and *Ph* models (fig. 5). In the *WhC* model, these percentages decreased to 7% and 6%, respectively (fig. 5). Similar findings were observed for presence/absence models (Fig. S26; Fig. S27).

**Figure 5.**
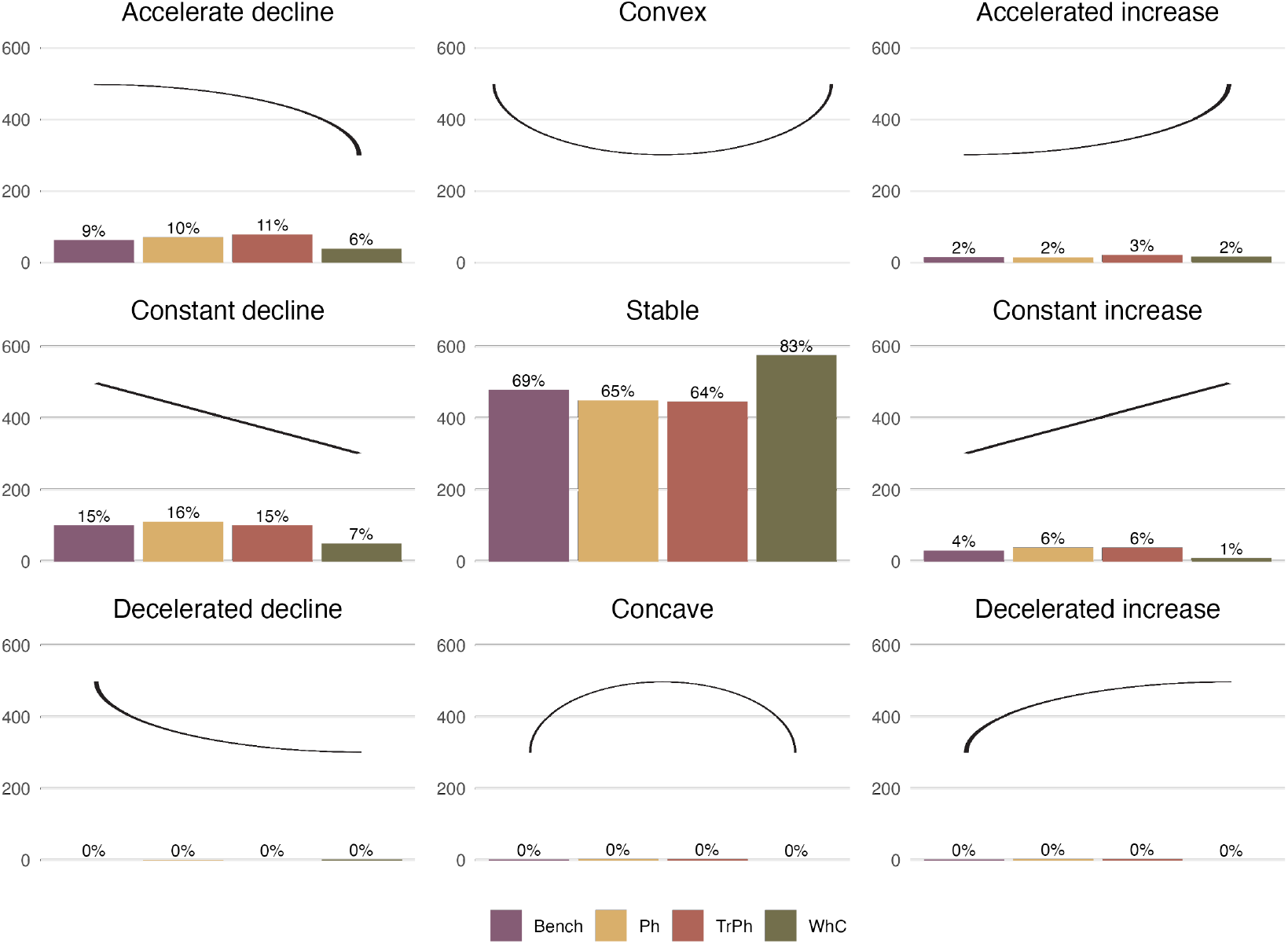
Number (y-axis) and proportion (computed across all coefficients for each model, and indicated above individual bars) of response curves (i.e. one for each species-predictor combination) following the typology (nine shapes highlighted by the black curve in each panel) of Rigal, Devictor, and Dakos (2020). Results are presented for the different model structures.

Both abundance and presence/absence *TrPh* models (which include species functional traits) reveal meaningful trait-environment relationships between the first fuzzy-PCA axis and the seven environmental predictors. This suggests that the occurrence of certain traits is favoured or hindered under certain environmental conditions (Fig. S28). For instance, mobile predatory species showed larger declines in abundance as fetch increases than sessile suspensivores (Fig. S28). Moreover, increase in organic matter concentration and decrease in current velocities were associated with higher abundances of suspensive feeders.

### Exploring the residual correlation

Residual species-species correlations were compared between the *Bench* model and the *WhC* model using both presence/absence (Fig S29) and abundance data (fig. 6). We only focused on this comparison because of the higher predictive performance and higher proportion of explained variance by random effects of the *WhC* model (fig. 4). Residual correlations estimated from both models were highly correlated, both for presence/absence and abundance data (fig. 6 and Fig. S29). However, agreement between models varied across different random effects from a moderate correlation between residuals associated with the Habitat random effect (R^2^ = 0.57) or with the Site random effect (R^2^ = 0.64), to a high correlation between residuals related to the Year random effect (R^2^ = 0.95). The δ index main modal distribution, which is centred on zero, confirms an overall agreement between residual correlations estimated from both models in relation to the Year random effect with a marginal proportion of sign changes (0.45% for correlations greater than |0.25|; fig. 6 B) only related to low species-species residual correlations (<0.25; fig. 6 A and Fig. S29). In contrast, the δ index highlights inconsistencies in both magnitude and sign changes between residuals associated with the Habitat and the Site (12.2% and 9.11% of sign changes related to correlation greater than |0.25|, respectively) random effects. Similar results were obtained for presence/absence models (Fig. S29).

**Figure 6.**
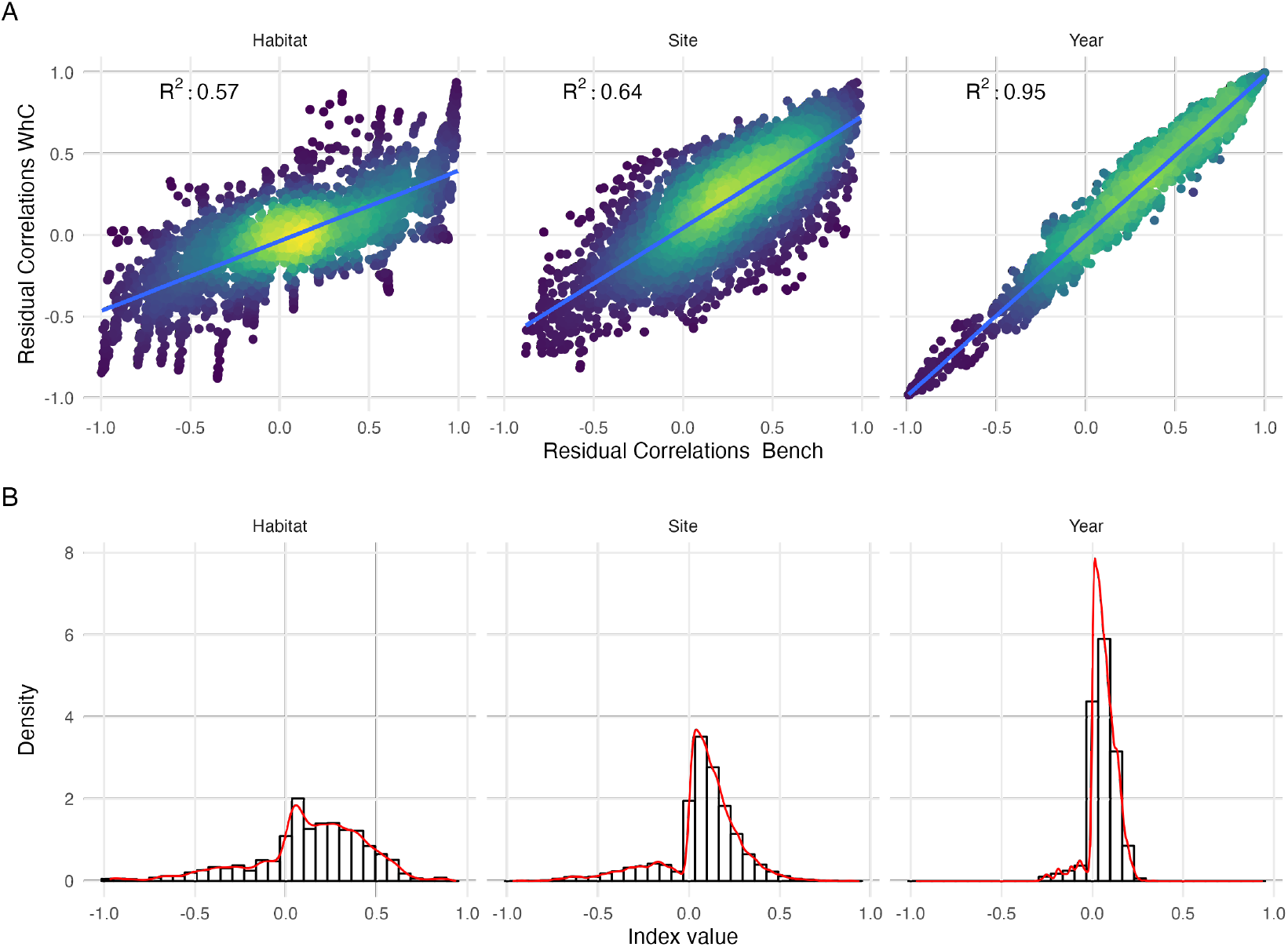
(A) Comparison of residual species-species correlations associated with the three random effects estimated by the *Whole Community Model* (*WhC*; y-axis) and the *Benchmark model* (*Bench*; x-axis) fitted on abundance data. The colour scale highlights the density of points in each scatter plot. (B) Distribution of the δ index characterising changes in sign (negative values indicate sign change) and magnitude (higher absolute values indicate higher numerical difference) between residual correlations estimated by the *WhC* model and the *Bench* model fitted with abundance data for the three random effects (Habitat, Site, Year).

## Discussion

Case studies in community ecology typically rely on partial and heterogeneous observations (Pollock, O’Connor, et al. 2020) but also on incomplete knowledge of target species ecological features (e.g. traits, phylogeny; Tyler et al. (2012)). This study investigated how *jSDM* performance varied depending on the type of information included (i.e. phylogeny, traits or data on non-target species) using a multi-assessment framework (spanning interpretability, inference and prediction, at both species- and community-level metrics, Table 1) enabling a thorough evaluation of model performance.

Using a typical case study, we found that although various *jSDM*s’ formulations could display decent and similar explanatory performances, their predictive power can be substantially lower and highlight larger differences between models. This discrepancy between explanatory and predictive powers may be due to the large number of rare species within our dataset. However, we found that *jSDM*’s performance, in particular the predictive power of abundance models at the species level, mostly increased when including information related to the 179 non-target species sampled alongside with the 99 polychaetes of interest. However, improvement in species-level predictions does not directly translate into enhanced performance at the community level. The *WhC* model did not improve estimates of beta diversity or total abundance relative to the other models and largely overpredicted species richness, as previously highlighted (Zurell, Pollock, and Thuiller 2018). Given *HMSC* hierarchical structure (Poggiato et al. 2021), inclusion of monitoring data related to other species likely improves model performance for the target assemblage by capturing relevant drivers that are not explicitly considered. For instance, it can help describe target species’ realised niches by accounting for ecological processes related to environmental conditions (including trait-mediated responses) or biotic interactions that are not explicitly captured otherwise (Ovaskainen, Tikhonov, Norberg, et al. 2017). In our case, the main differences between residual correlations estimated by the *Benchmark model* and the Whole community model relate to spatial random effects (i.e. site and habitat). In contrast, the year random effect yielded similar residual co-occurrences in both models. This suggests that including non-target species in our case, mostly helped capture spatial variability in species associations across sites and habitats.

*jSDM*s have already been used to model the distribution of a wide variety of species ranging from micro-organisms (Minard et al. 2019) to megafauna (Brimacombe, Bodner, and Fortin 2021) inhabiting many different ecosystems. Here, while we studied assemblages associated with two specific coastal habitats, i.e. seagrass and sand, that have original characteristics as they are located at the land-sea interface (Boyé, Thiébaut, et al. 2019), our case study reflects typical aspects of applied ecological research. These include issues related to data limitations and availability but also typical features of ecological communities (e.g. prevalence of rare and transient species; Magurran and Henderson (2003); Snell Taylor et al. (2018)). Valuable insights on trait-environment relationships are scarce in our study, potentially indicating that the contribution of functional ecology to *jSDM*s can be limited by trait data quality and availability (Tyler et al. 2012; Juan et al. 2022). For instance, we found an interaction between trophic modalities (i.e. microphagous versus macrophagous diet) and fetch (Fig. S15), indicating that organisms that filter on small particles are less likely to occur in wave-exposed sites where high levels of sediment resuspension can block their filtering systems (Manning, Peterson, and Bishop 2014). Yet, the limited number of informative trait-environment relationships or species-environment relationships either suggest that neutral processes may shape polychaete assemblages (Boyé, Thiébaut, et al. 2019) or highlight a mismatch between trait data, environmental data, and the ecological processes at play (Juan et al. 2022). For instance, the environmental variables we have used only capture average climatic conditions, but do not allow quantifying the environmental variability prevailing in the coastal environment, such as extreme events, seasonal or annual variability. Likewise, the list of available fuzzy-coded traits only partially captures species capacity to adapt to environmental variability (Juan et al. 2022). Thus, effectiveness of inclusion of traits in *jSDM*s is likely to be limited, or to rely on effort to collect relevant trait information. In our case, while including traits does not improve model predictive power, it somehow enhances our understanding of species responses along environmental gradients. Hence, if the goal is not prediction but inference (Tredennick et al. 2021), including traits and proxies of phylogeny can facilitate *jSDM* interpretation.

Importantly, while we show that including non-target species can improve predictive performance, this benefit might vary depending on the robustness of non-target species monitoring data (e.g. detection issues), their role within the ecosystem (e.g. engineer species are likely more influential in local communities than rare transient species), or processes shaping the target assemblage (i.e. if the influence of abiotic factors dominates, then adding other species will likely have marginal consequences on model performance). While the list of additional species to consider can be prioritised based on existing knowledge of well-studied ecosystems, such information is often unavailable. Further studies, conducted on other ecosystems or using simulated datasets to overcome limitations related to real world datasets (DiRenzo, Hanks, and Miller 2022), could help determine specific criteria, such as the optimal number of non-target species to include to improve models’ performance. While species communities and assemblages are largely defined arbitrarily (Stroud et al. 2015), a systematic assessment of *jSDM* performance in response to variations in the number and types (for instance based on their functional or trophic roles) of non-target species would be valuable to optimise model performance.

This paper lays out an original framework to systematically compare multiple facets of alternative *jSDM* formulations (i.e. including phylogeny, traits or additional species) on model interpretability, explanatory and predictive power (Table 1). Using a set of complementary metrics, we specifically assess performance of alternative model formulations fitted to presence-absence or abundance data at the species and community levels. Our framework goes beyond existing guidelines proposed to assess the performance of *jSDM* fitted on presence-absence data (Wilkinson et al. 2021) or that focus on the predictive power of abundance-based models (e.g. Waldock et al. (2022)). It specifically compares the performance (both explanatory and predictive) and interpretability of alternative models’ formulations accounting for the multiple and high-dimensional components that are typical of *jSDM*s, namely (1) species and community level predictions including alpha and beta diversity metrics and ranking of predictions according to species prevalence/abundance; (2) species-environment relationships where we transposed the framework initially developed for time series by Rigal, Devictor, and Dakos (2020) into an effective tool to classify response curves according to 9 categories; (3) trait-environment relationships; and (4) residual species-species correlations associated with random effects thanks to a new index that allows capturing changes in the sign and the magnitude of residual correlations.

Overall, our results provide new insights into the most appropriate strategies for *jSDM* fitting, according to modelling objectives (Tredennick et al. 2021) and available data. While the four models considered had similar explanatory power, adding extra information to standard *jSDM*s that only consider abiotic predictors can prove useful in some cases. For instance, adding monitoring data on other non-target species can substantially increase a model’s predictive power by modifying inferred species-environment relationships and residual correlation matrices. Similarly, adding traits or phylogeny can improve a model’s interpretability. Future studies will be key to consolidate our findings on simulated case studies (Zurell, Berger, et al. 2010; DiRenzo, Hanks, and Miller 2022), or across contrasted ecosystems, for instance dominated either by environmental filtering, or by competitive processes. Generalising this approach across ecosystems will further help prioritise data collection effort in the long term. For this purpose, we recommend using a multi-model inference framework similar to the one used in this study to systematically assess trade-offs associated with alternative *jSDM*s formulations.

## Author Contributions

MPM conceived the project with inputs from CV, AB, MC. CV analysed data and wrote the first draft. All authors provided significant inputs and approved the final version of the manuscript.

## Supporting information

Supplementary results

## Acknowledgments

We are grateful to Marion Maguer and Vincent Le Garrec who conducted fieldwork and laboratory analyses, as well as to supporting students and staff involved in the REBENT monitoring programme coordinated by Sandrine Derrien (MNHN) and its funding partners (Agence de l’eau Loire-Bretagne, Région Bretagne, DREAL Bretagne). The authors would also like to acknowledge the Pôle de Calcul et de Données Marines (PCDM) for providing DATARMOR storage and computational resources. https://pcdm.ifremer.fr. MPM is the recipient of an ANR early career grant ANR-21-CE02-0006.

