## Supplementary results for "A Multifaceted Framework To Assess tradeoffs in Interpretability, Explanatory and Predictive Performances Of Alternative Joint Species Distribution Models"

### Appendix A - Data Sources and Descriptions of the Datasets

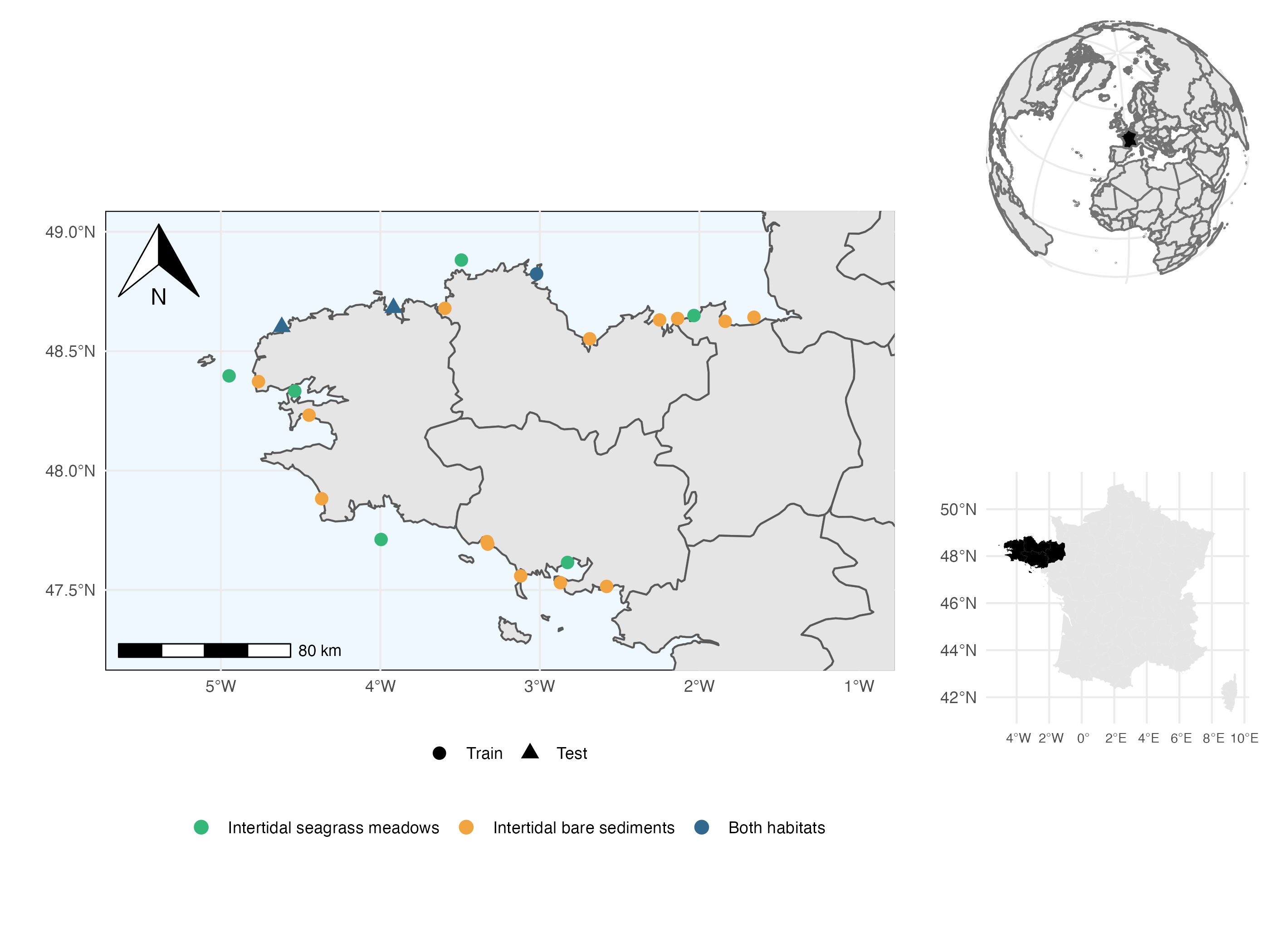

Figure 1: Map of the sampled sites. Point shapes vary according to whether they belong to the training set (circles ; used to evaluate model explanatory power) or the test set (triangles ; used to evaluate model predictive power). Point colours vary according to the presence or absence of the two habitats in each site. The two test sites include the two habitats (i.e. seagrass and bare sand) and were chosen because they occur in average environmental conditions at the scale of the region (thereby limiting extrapolation of the model) but still harbour different communities, representative of the known diversity gradient across the region.

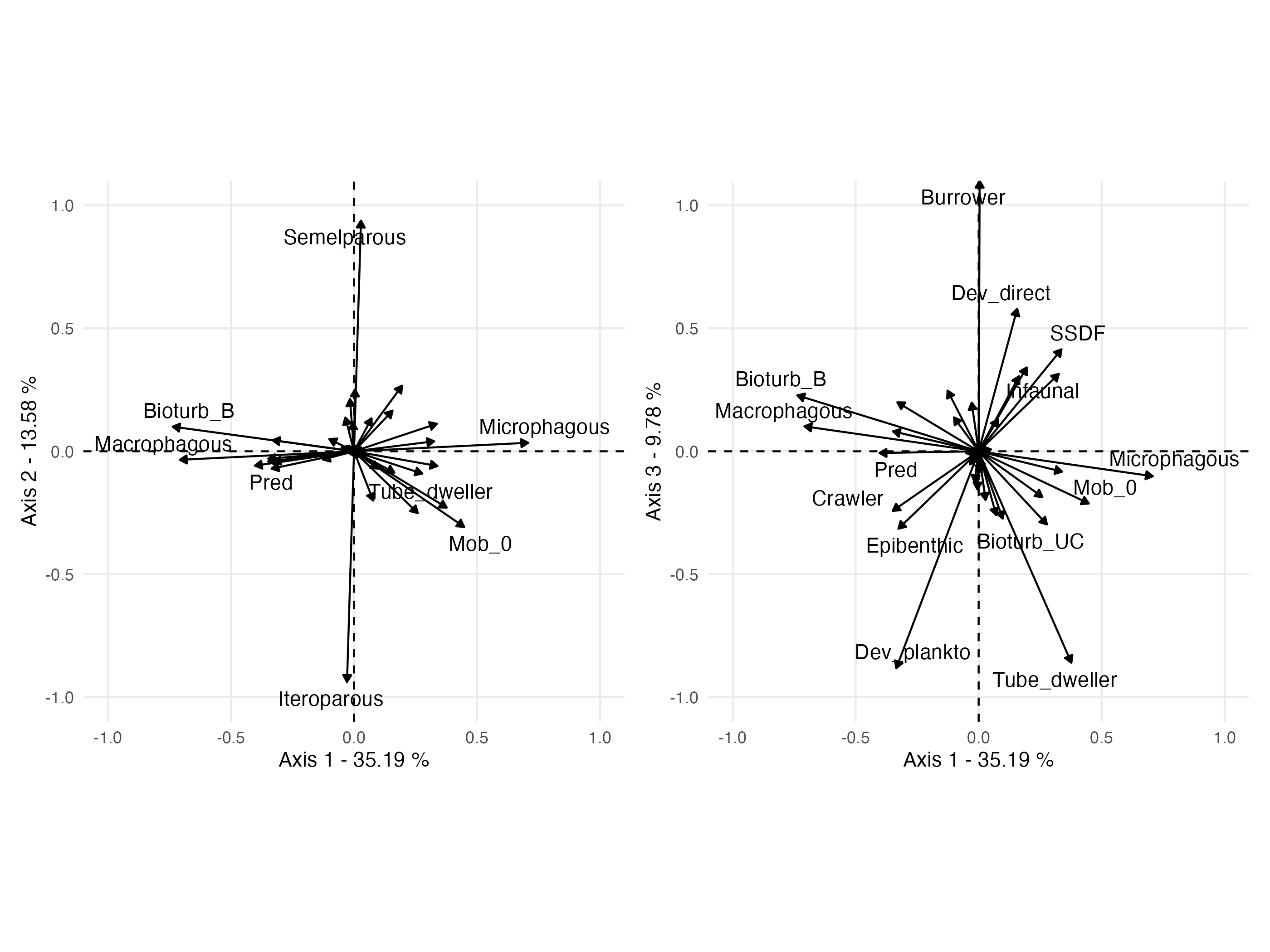

Figure 2: Fuzzy PCA of the species-by-trait matrix. The first three axes represent 58.55% of the total variance. The first axis distinguishes sessile microphagous species (top positive values) from mobile macrophagous predatory species (bottom negative values). The second axis is a gradient of reproductive strategies (semelparous vs iteroparous). The third axis distinguishes burrowers with direct development from tube-dwellers with planktonic development. For abbreviations and meaning of the trait modalities, see Boyé *et al.* (2019).

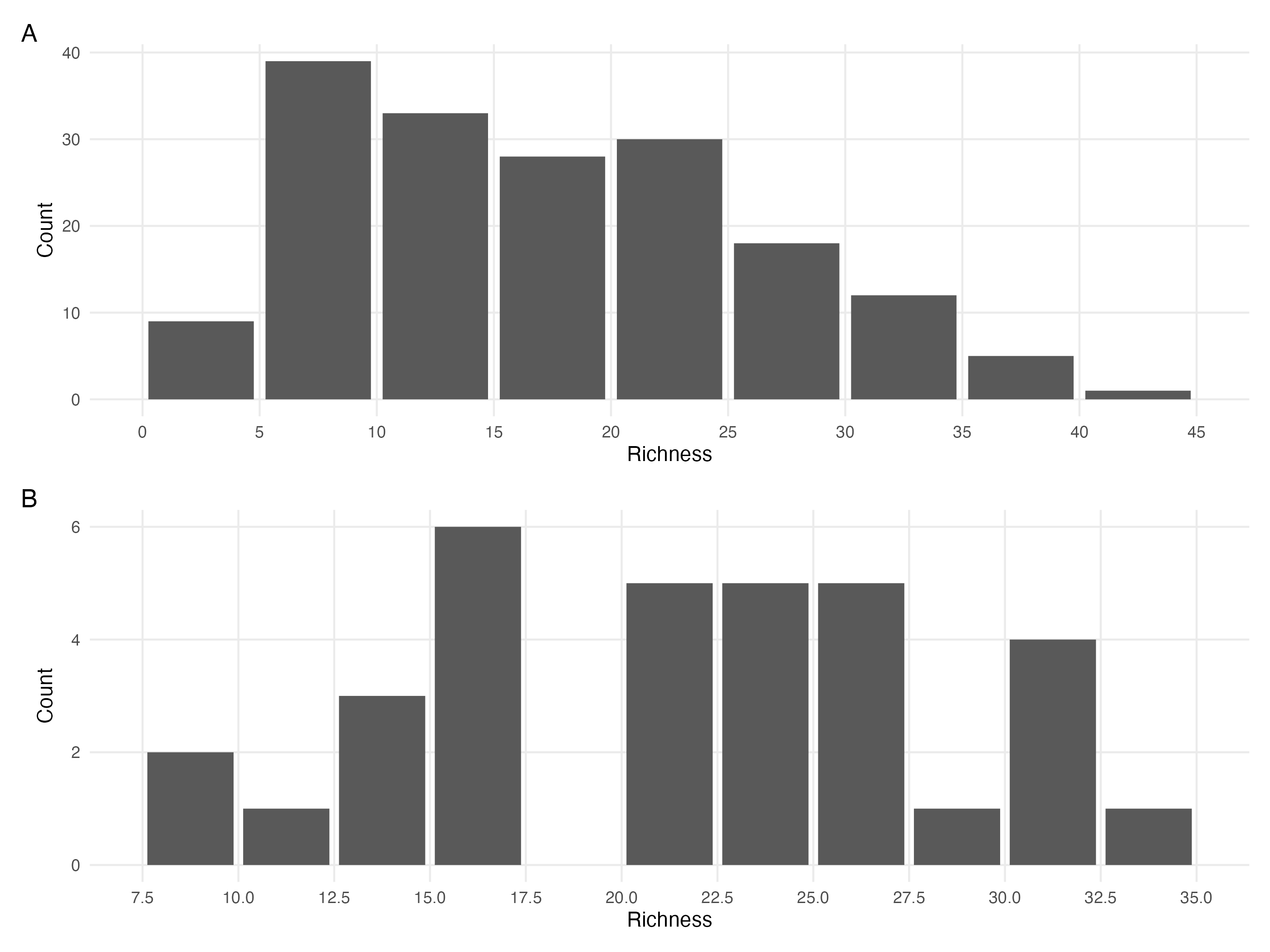

Figure 3: A. Distribution of the number of species in the samples (site times habitat times year) of the train dataset. B. Same distribution for the test dataset.

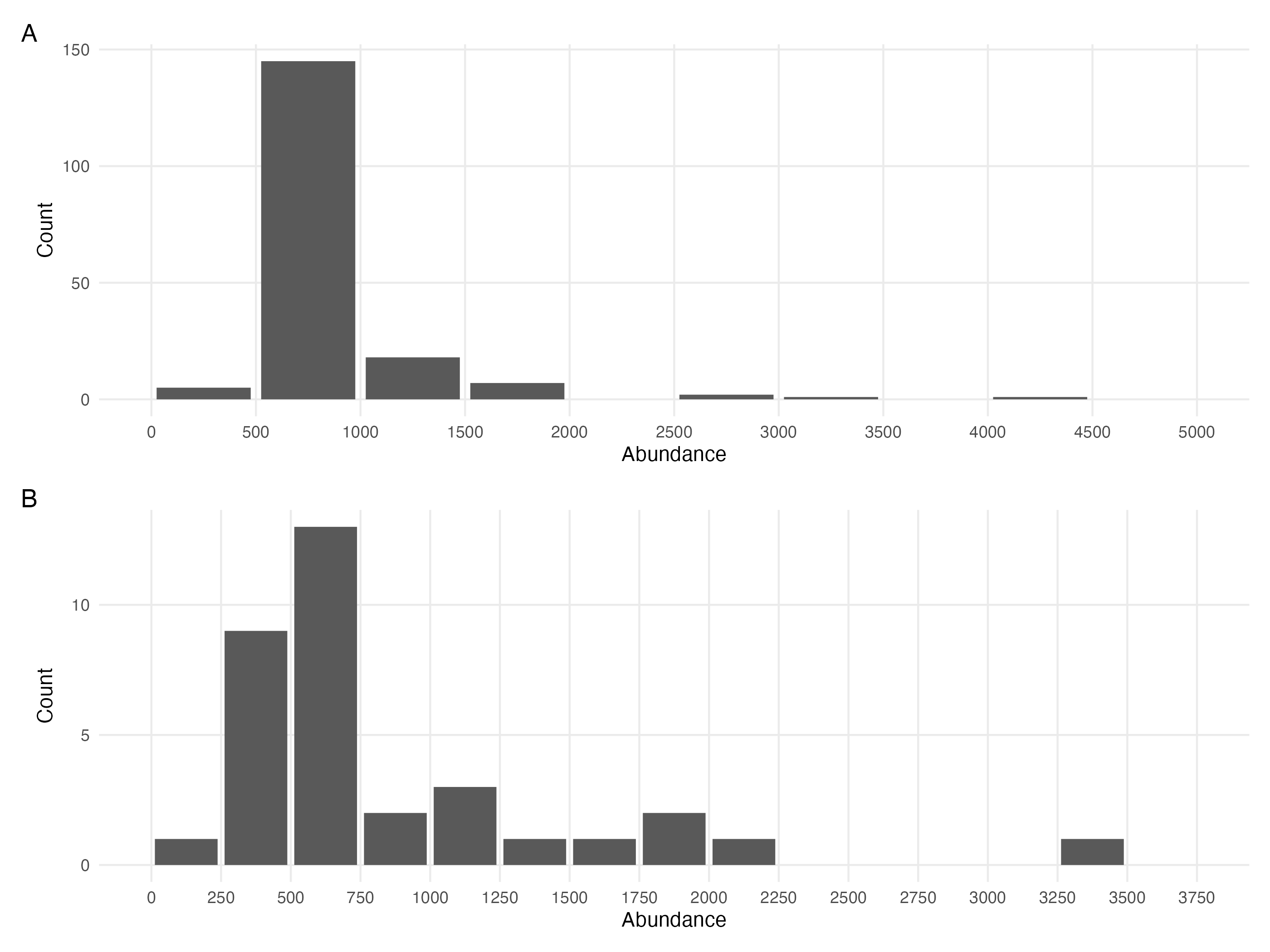

Figure 4: A. Distribution of the total number of individuals (abundance) in the samples (site x habitat x year) of the train dataset. B. Same distribution for the test dataset.

#### Environmental data acquisition

The models used in this study were fitted with predictors consisting of a set of seven environmental variables related to oceanography, hydrography, and granulometry obtained from Boyé (2019) The oceanographic variables include the standard deviation of salinity, surface water temperature, mean current velocity, and fetch, which were obtained from the PREVIMER database (Lecornu & Roeck 2009) based on the MARS3D model (Lazure & Dumas 2008). The variables were averaged by extracting daily data for the sampled year at the site coordinates with a buffer including the eight adjacent cells. The fetch was calculated as the average length of nine radiating fetch segments with a maximum distance of 300km. The granulometry variables were derived from sediment cores that were taken along with the associated fauna. The cores were dried, separated into 15 fractions, and the Trask index was calculated as the ratio of the 25th to 75th percentile of the grain distribution. Organic matter mass was estimated through the loss of mass after combustion in an oven.

### Appendix B - Model Convergence

#### Environmental coefficients

Table 1: Potential scale reduction factors (PSRF) and effective sample sizes (ESS) for environmental regression parameters (i.e beta coefficients) estimated for the four different models (Bench, Ph, TrPh, WhC) fitted either to abundance or presence-absence data. For further details see Fig. S5 to Fig. S12.

| Model | Data Type | Number of coefficients | PSRF (mean $\pm$ sd) | ESS (mean $\pm$ sd) |
| --- | --- | --- | --- | --- |
| Benchmark | Abundance | 1485 | 1.18 $\pm$ 0.267 | 701 $\pm$ 576 |
| Benchmark | Presence/Absence | 1485 | 1.00 $\pm$ 0.002 | 4967 $\pm$ 417 |
| Phylogeny | Abundance | 1485 | 1.18 $\pm$ 0.204 | 566 $\pm$ 420 |
| Phylogeny | Presence/Absence | 1485 | 1.00 $\pm$ 0.001 | 4947 $\pm$ 408 |
| Traits & Phylogeny | Abundance | 1485 | 1.21 $\pm$ 0.317 | 489 $\pm$ 358 |
| Traits & Phylogeny | Presence/Absence | 1485 | 1.00 $\pm$ 0.008 | 11459 $\pm$ 2649 |
| Whole Community | Abundance | 4170 | 1.21 $\pm$ 0.287 | 739 $\pm$ 631 |
| Whole Community | Presence/Absence | 4170 | 1.00 $\pm$ 0.002 | 4962 $\pm$ 406 |

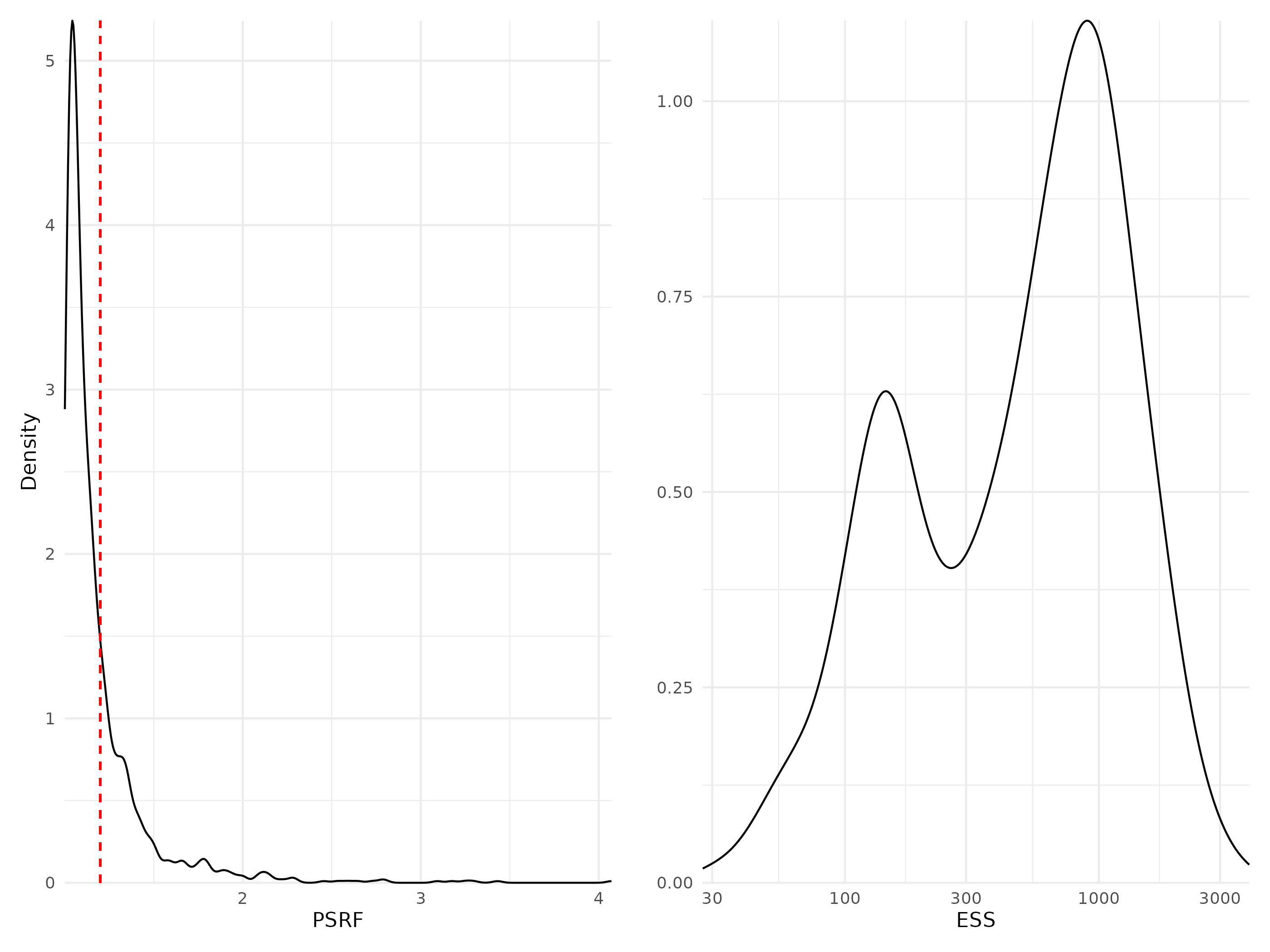

Figure 5: Density curves of potential scale reduction factors (PSRF see Brooks & Gelman (1998); left panel) and effective sample sizes (ESS; right panel) for Beta regression parameters (i.e environmental coefficients) estimated for the benchmark model fitted with abundance data. For PSRF, values greater than 1.2 (dotted red line) indicate potential convergence issues. ESS estimates the number of independent samples used to estimate each parameter (the more the better).

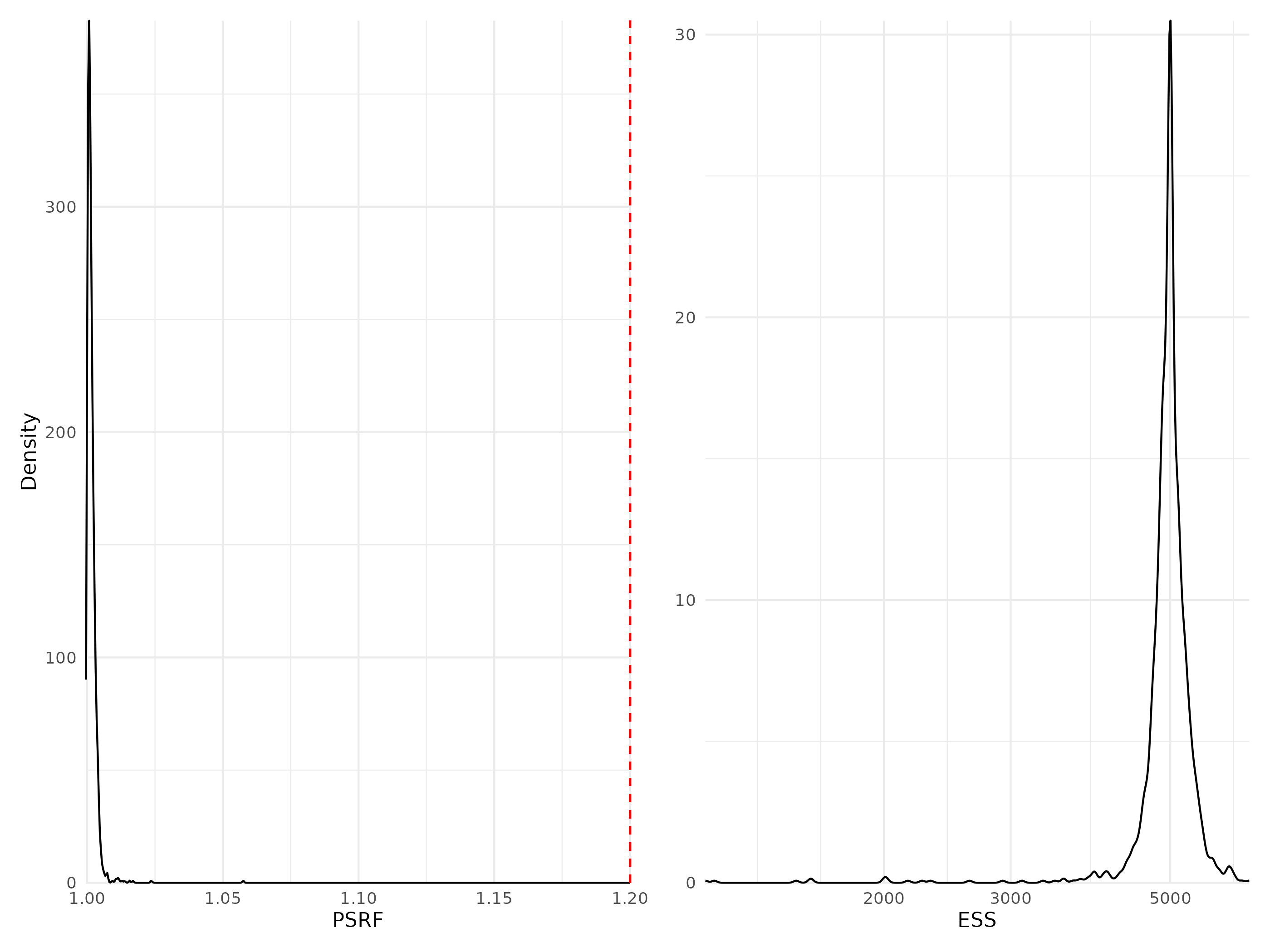

Figure 6: Density curves of potential scale reduction factors (PSRF see Brooks & Gelman (1998); left panel) and effective sample sizes (ESS; right panel) for Beta regression parameters (i.e environmental coefficients) estimated for the benchmark model fitted with presence/absence data. For further details see Fig. S5.

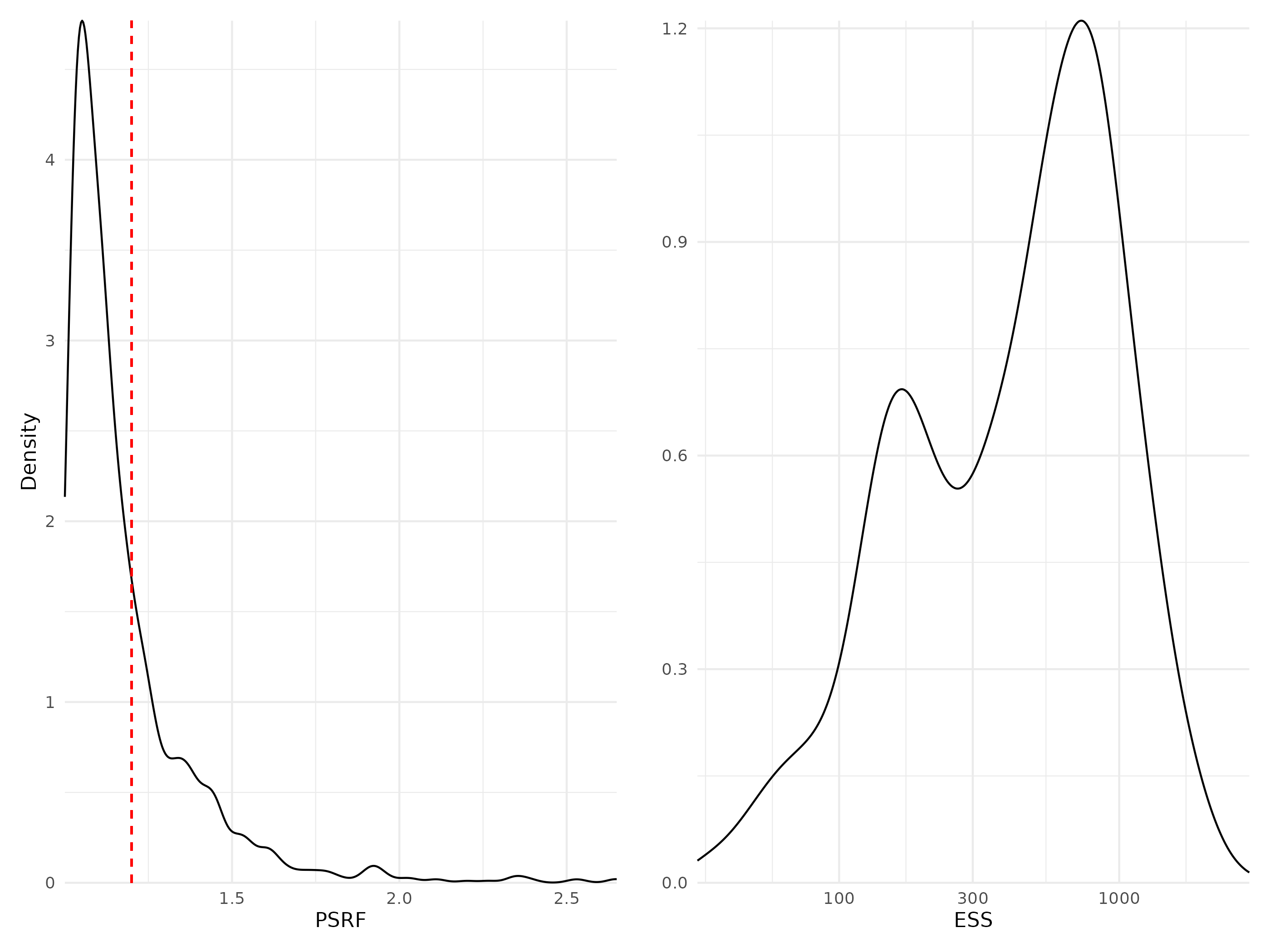

Figure 7: Density curves of potential scale reduction factors (PSRF see Brooks & Gelman (1998); left panel) and effective sample sizes (ESS; right panel) for Beta regression parameters (i.e environmental coefficients) estimated for the phylogeny model fitted with abundance data. For further details see Fig. S5.

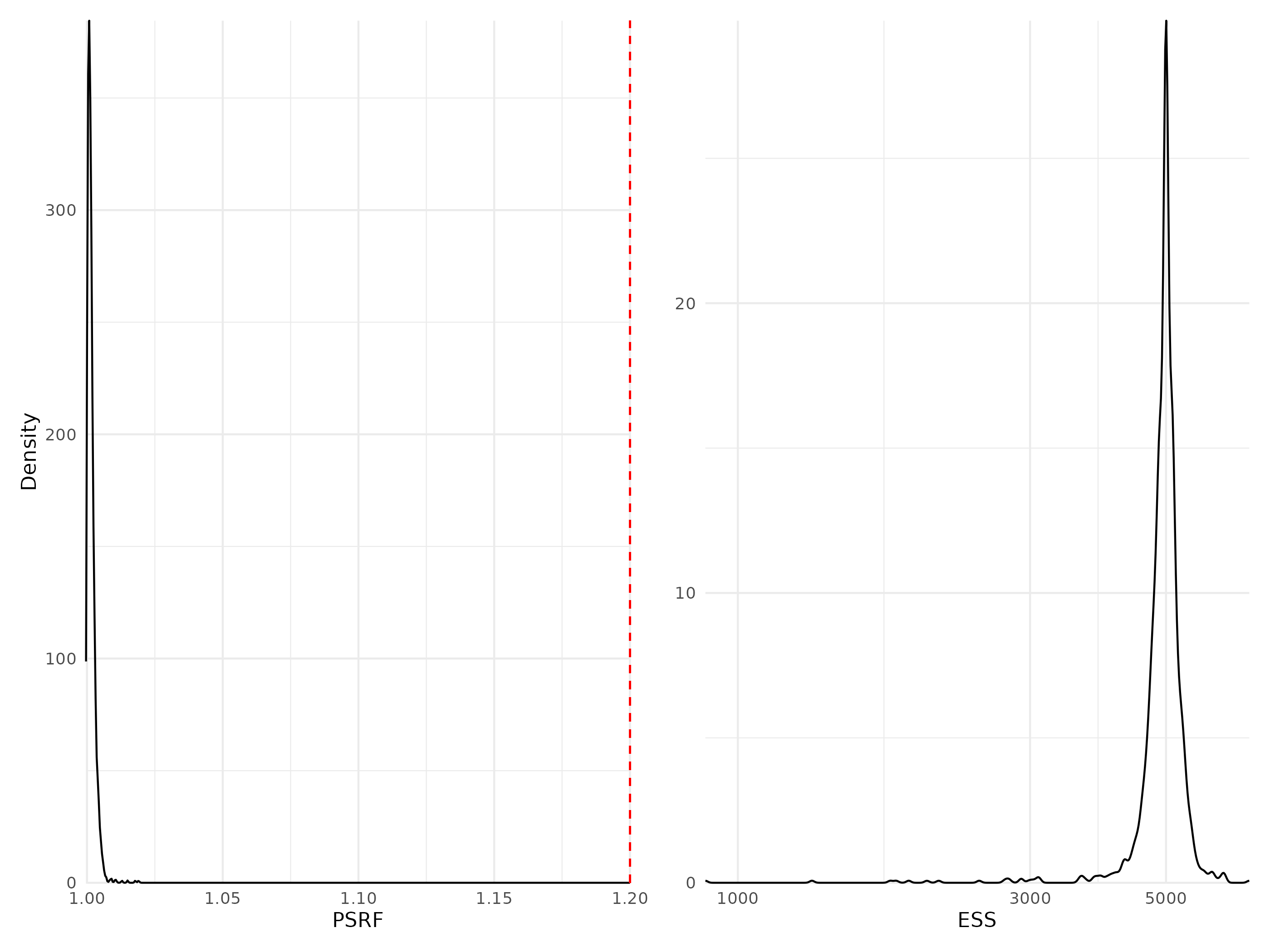

Figure 8: Density curves of potential scale reduction factors (PSRF see Brooks & Gelman (1998); left panel) and effective sample sizes (ESS; right panel) for Beta regression parameters (i.e environmental coefficients) estimated for the phylogeny model fitted with presence/absence data. For further details see Fig. S5.

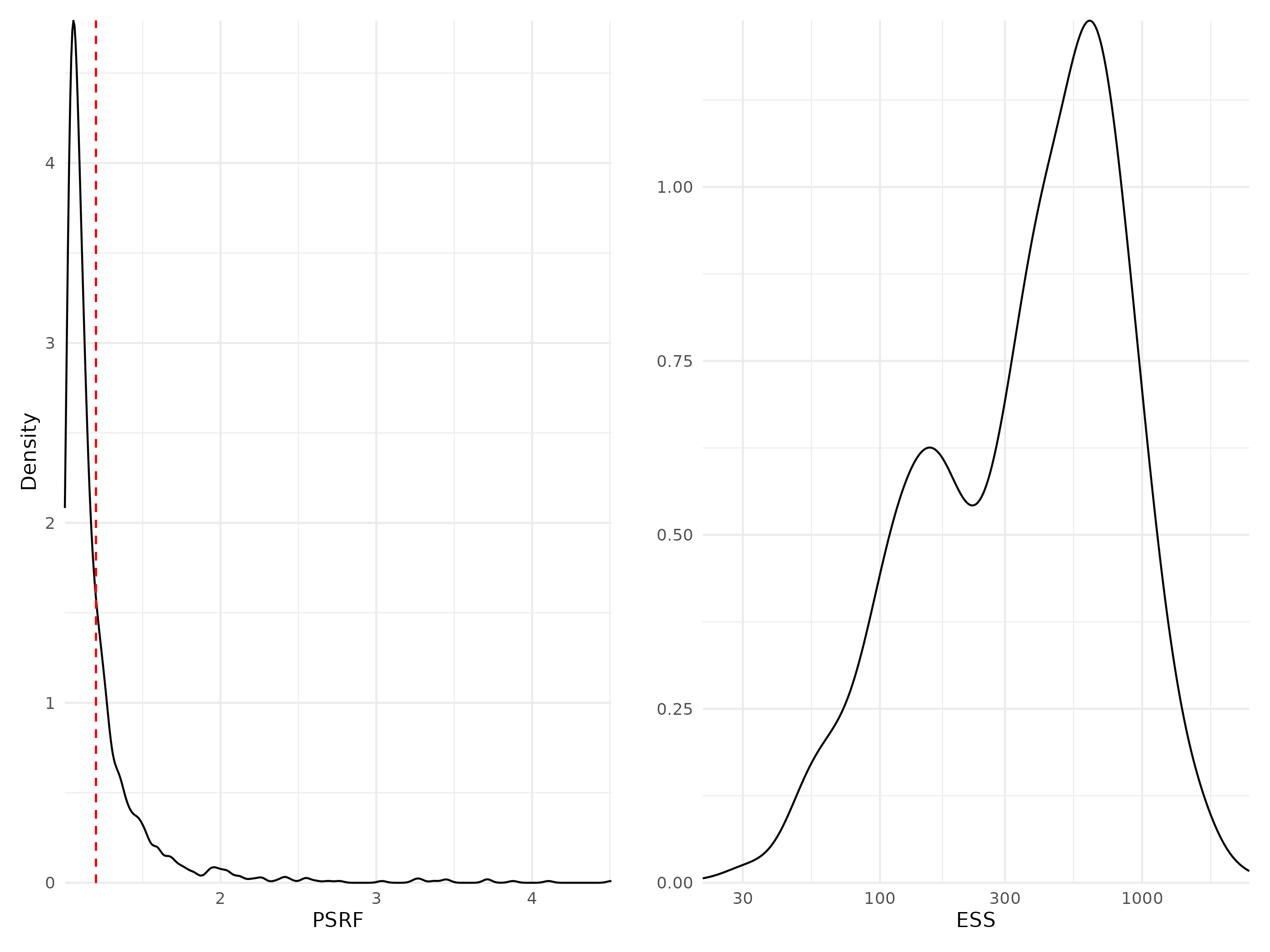

Figure 9: Density curves of potential scale reduction factors (PSRF see Brooks & Gelman (1998); left panel) and effective sample sizes (ESS; right panel) for Beta regression parameters (i.e environmental coefficients) estimated for the traits & phylogeny model fitted with abundance data. For further details see Fig. S5.

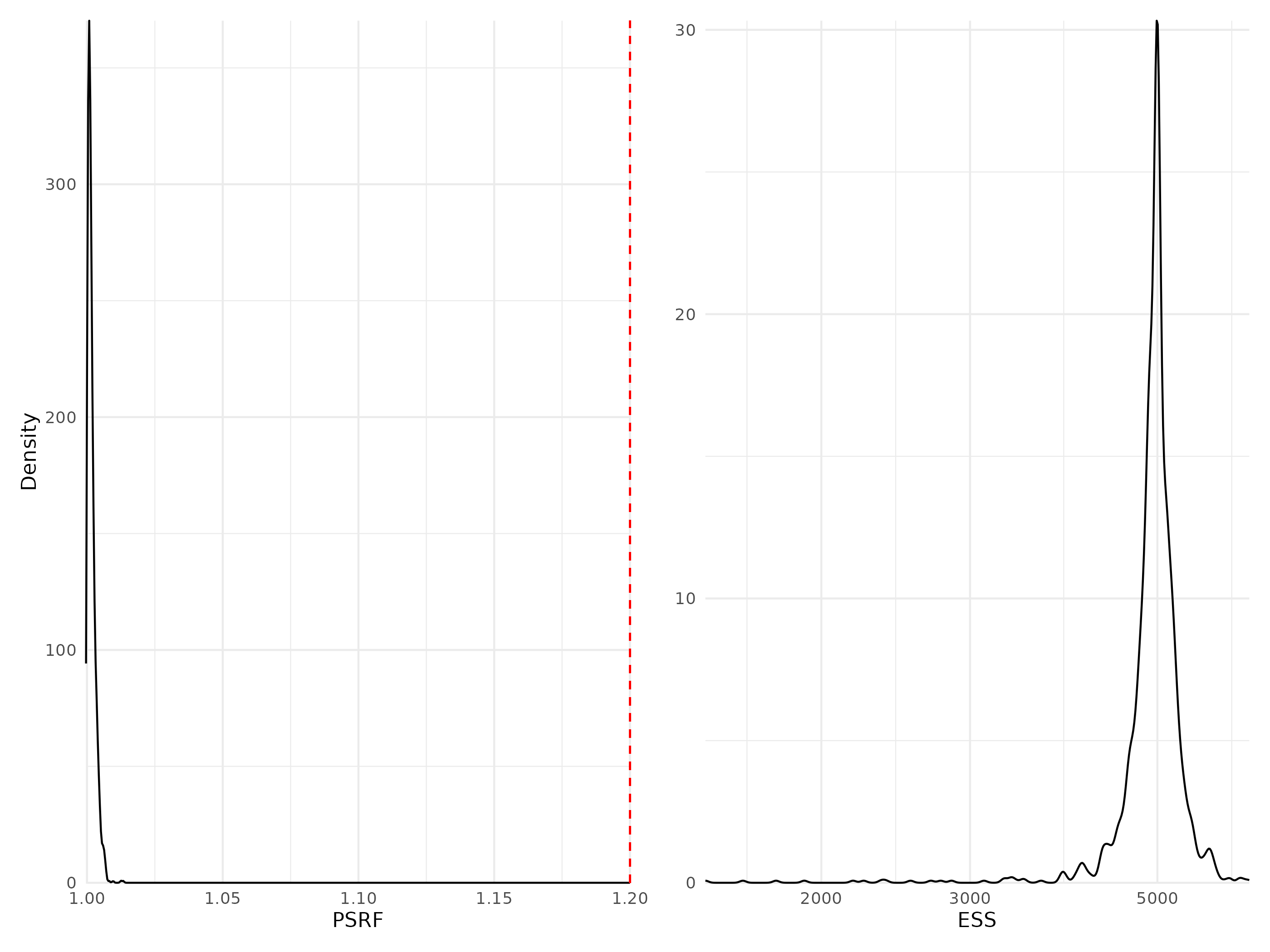

Figure 10: Density curves of potential scale reduction factors (PSRF see Brooks & Gelman (1998); left panel) and effective sample sizes (ESS; right panel) for Beta regression parameters (i.e environmental coefficients) estimated for the traits & phylogeny model fitted with presence/absence data. For further details see Fig. S5.

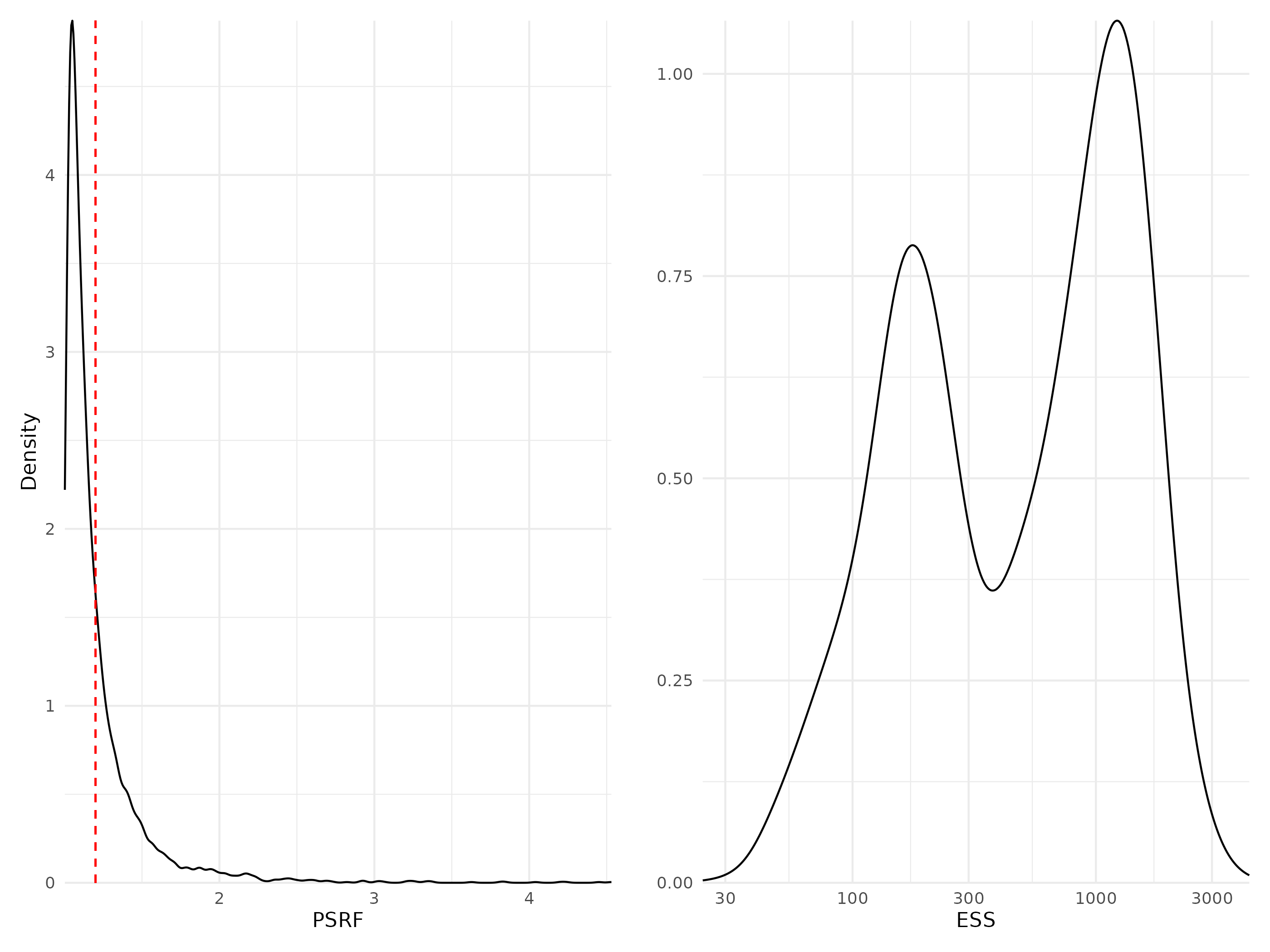

Figure 11: Density curves of potential scale reduction factors (PSRF see Brooks & Gelman (1998); left panel) and effective sample sizes (ESS; right panel) for Beta regression parameters (i.e environmental coefficients) estimated for the whole community model fitted with abundance data. For further details see Fig. S5.

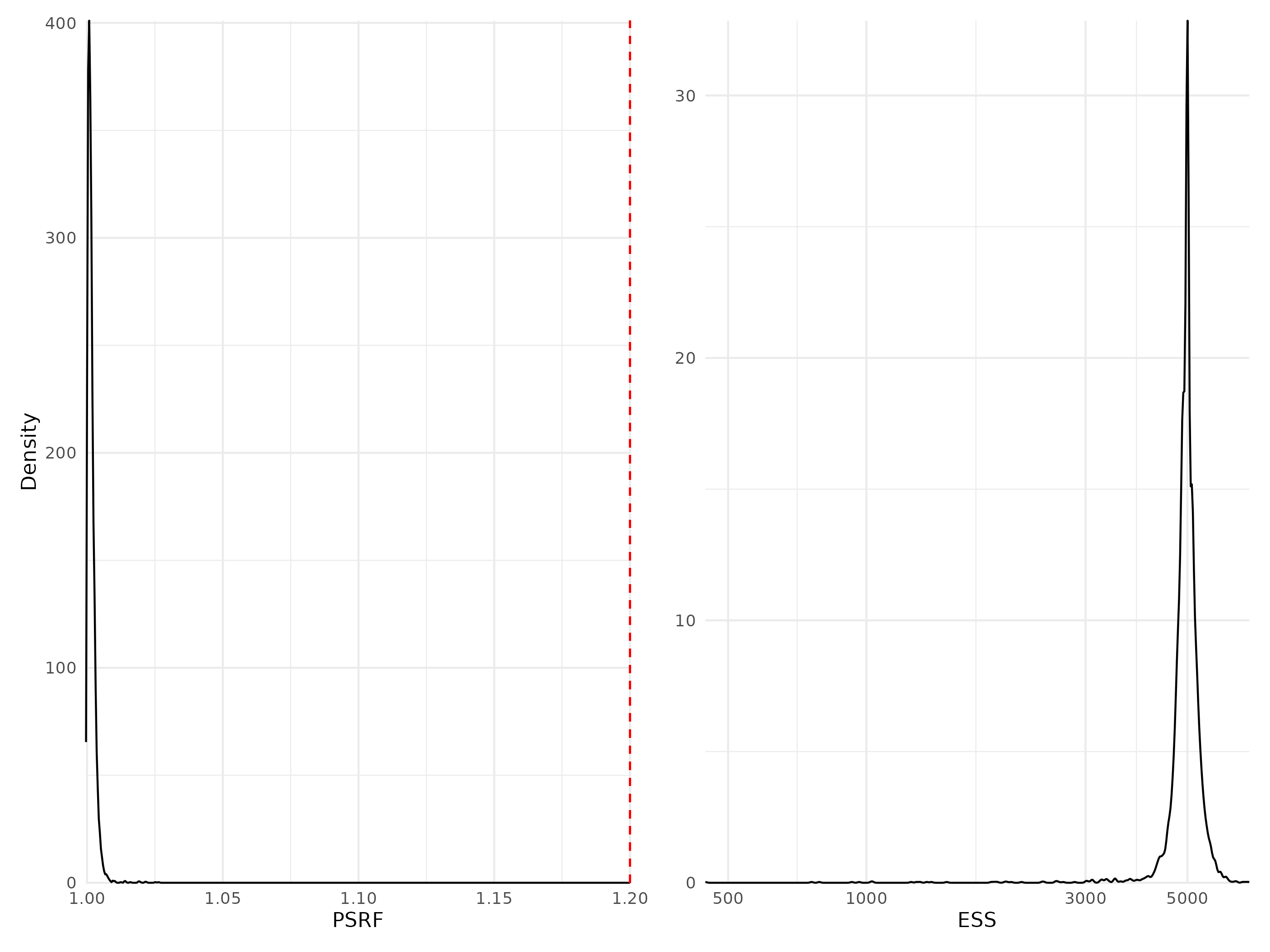

Figure 12: Density curves of potential scale reduction factors (PSRF see Brooks & Gelman (1998); left panel) and effective sample sizes (ESS; right panel) for Beta regression parameters (i.e environmental coefficients) estimated for the whole community model fitted with presence/absence data. For further details see Fig. S5.

#### Traits coefficients

Table 2: Potential scale reduction factors (PSRF) and effective sample sizes (ESS) for trait-environment regression parameters (i.e gamma coefficients) estimated for the model including trait information fitted either to abundance or presence-absence data. For further details see Fig. S13 to Fig. S14.

| Model | Data type | Number of coefficients | PSRF (mean $\pm$ sd) | ESS (mean $\pm$ sd) |
| --- | --- | --- | --- | --- |
| Traits & Phylogeny | Abundance | 60 | 1.08 $\pm$ 0.092 | 1232 $\pm$ 1209 |
| Traits & Phylogeny | Presence/Absence | 60 | 1.00 $\pm$ 0.001 | 13227 $\pm$ 1897 |

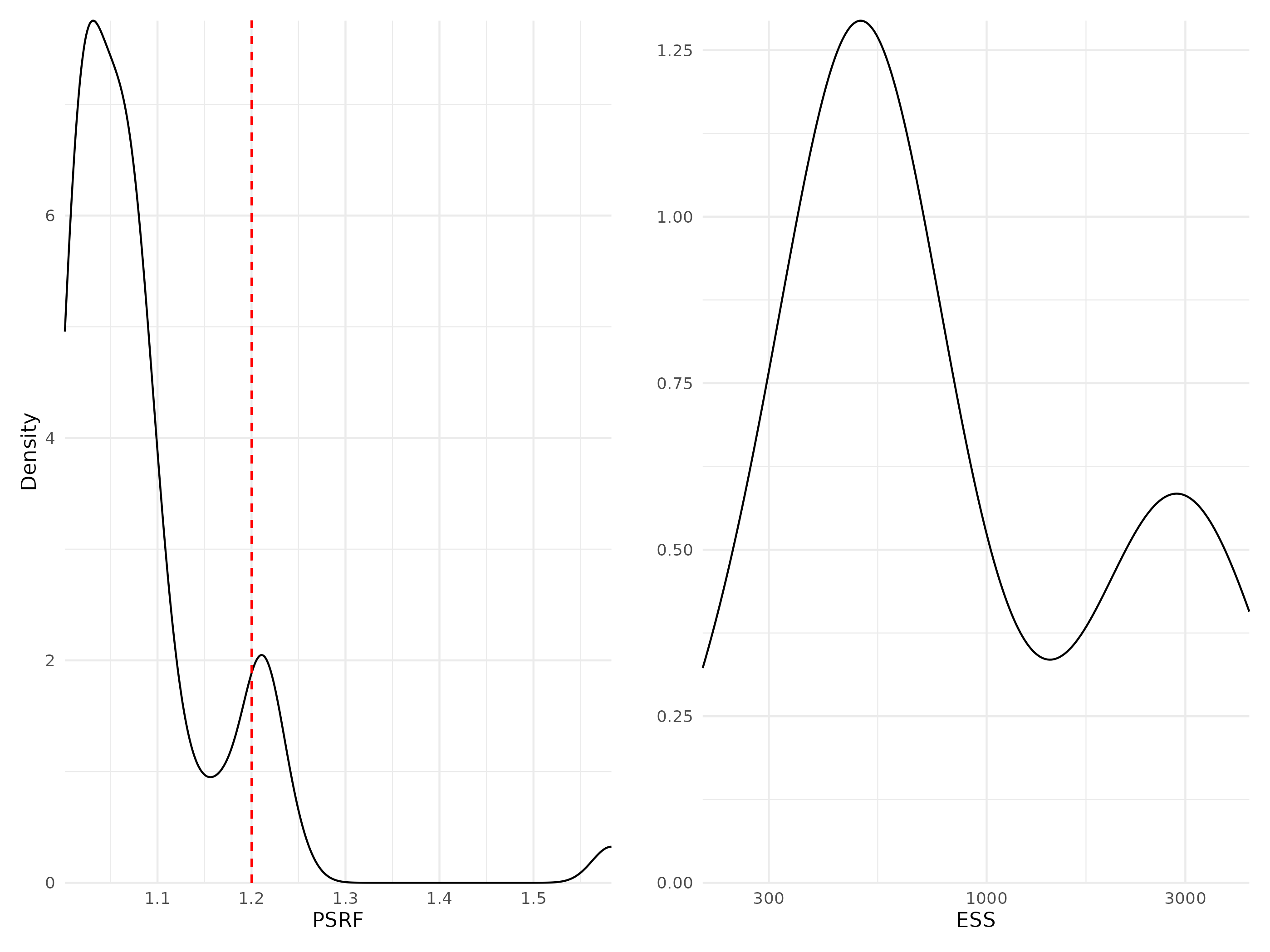

Figure 13: Density curves of potential scale reduction factors (PSRF see Brooks & Gelman (1998); left panel) and effective sample sizes (ESS; right panel) for Gamma regression parameters (i.e coefficients associated with trait-environment relationships, modeling how species traits influence their niches) estimated for the traits & phylogeny model fitted with abundance data. For further details see Fig. S5.

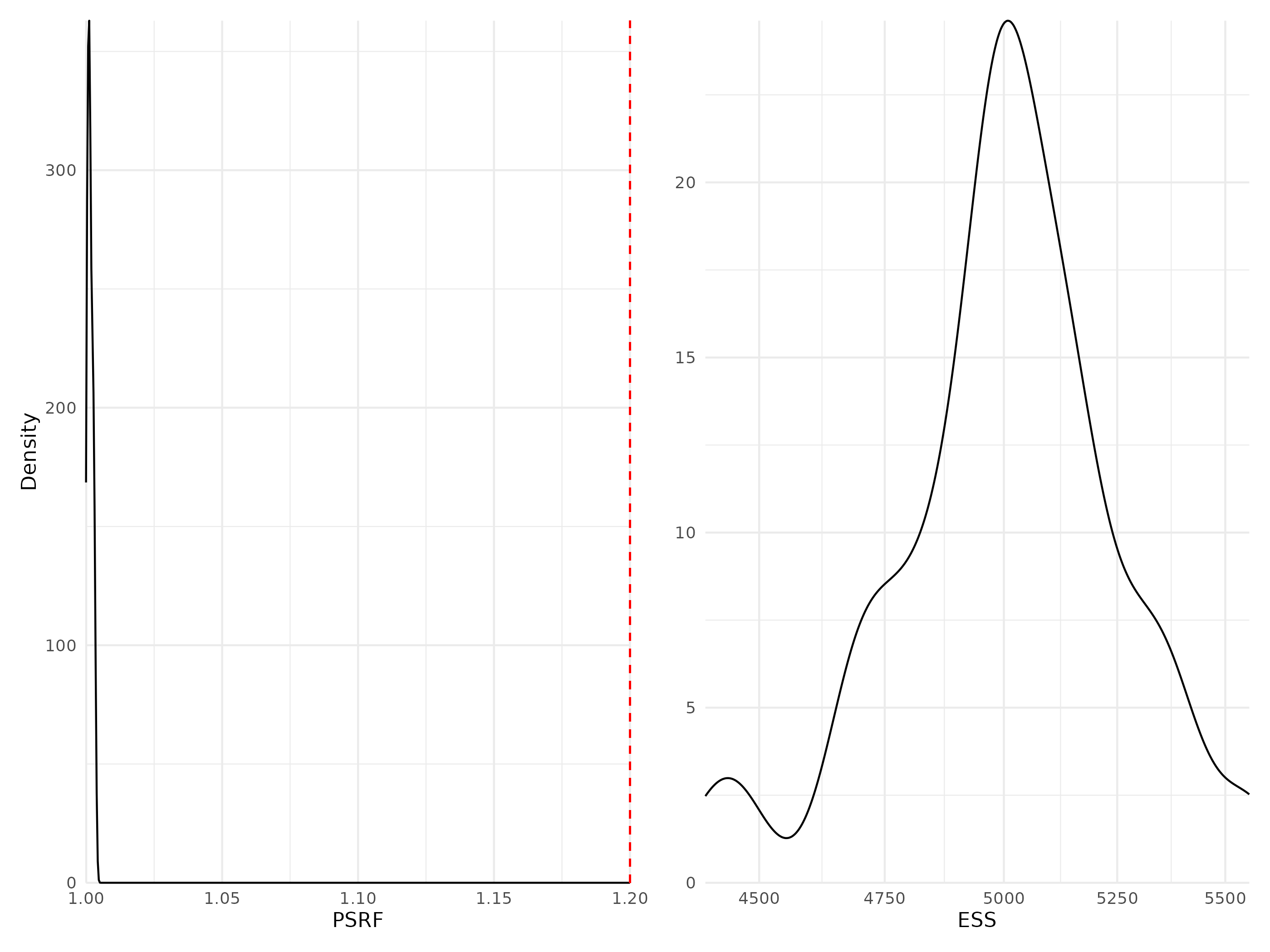

Figure 14: Density curves of potential scale reduction factors (PSRF see Brooks & Gelman (1998); left panel) and effective sample sizes (ESS; right panel) for Gamma regression parameters (i.e coefficients associated with trait-environment relationships, modeling how species traits influence their niches) estimated for the traits & phylogeny model fitted with presence/absence data. For further details see Fig. S5.

#### Phylogeny coefficients

Table 3: Potential scale reduction factors (PSRF) and effective sample sizes (ESS) for rho regression parameters (i.e phylogeny coefficient) estimated for the two models including phylogenetic information (TrPh and Ph).

| Model | Data type | Number of coefficients | PSRF | ESS |
| --- | --- | --- | --- | --- |
| Phylogeny | Abundance | 1 | 1.07 | 649 |
| Phylogeny | Presence/Absence | 1 | 1.00 | 9349 |
| Traits & Phylogeny | Abundance | 1 | 1.15 | 757 |
| Traits & Phylogeny | Presence/Absence | 1 | 1.00 | 5000 |

#### Link between model convergence and species response curves

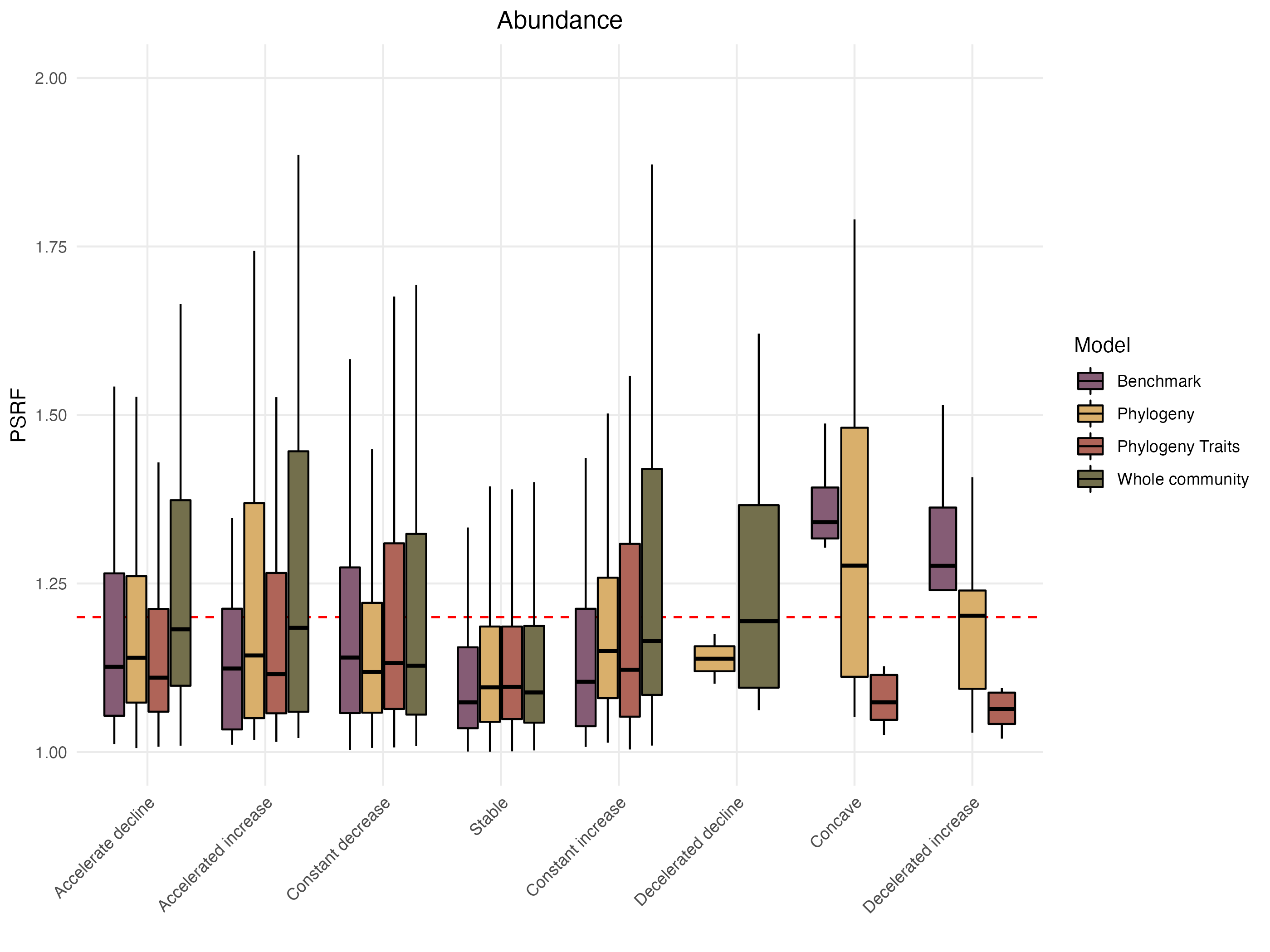

Figure 15: Distribution of the Potential scale reduction factors (PSRF) as a function of the different shapes of the response curves classified following the methodology proposed by Rigal *et al.* (2020) (see section “Assessing model performance and interpretability” for more details on the calculation methodology). Results for models fitted with abundance data.

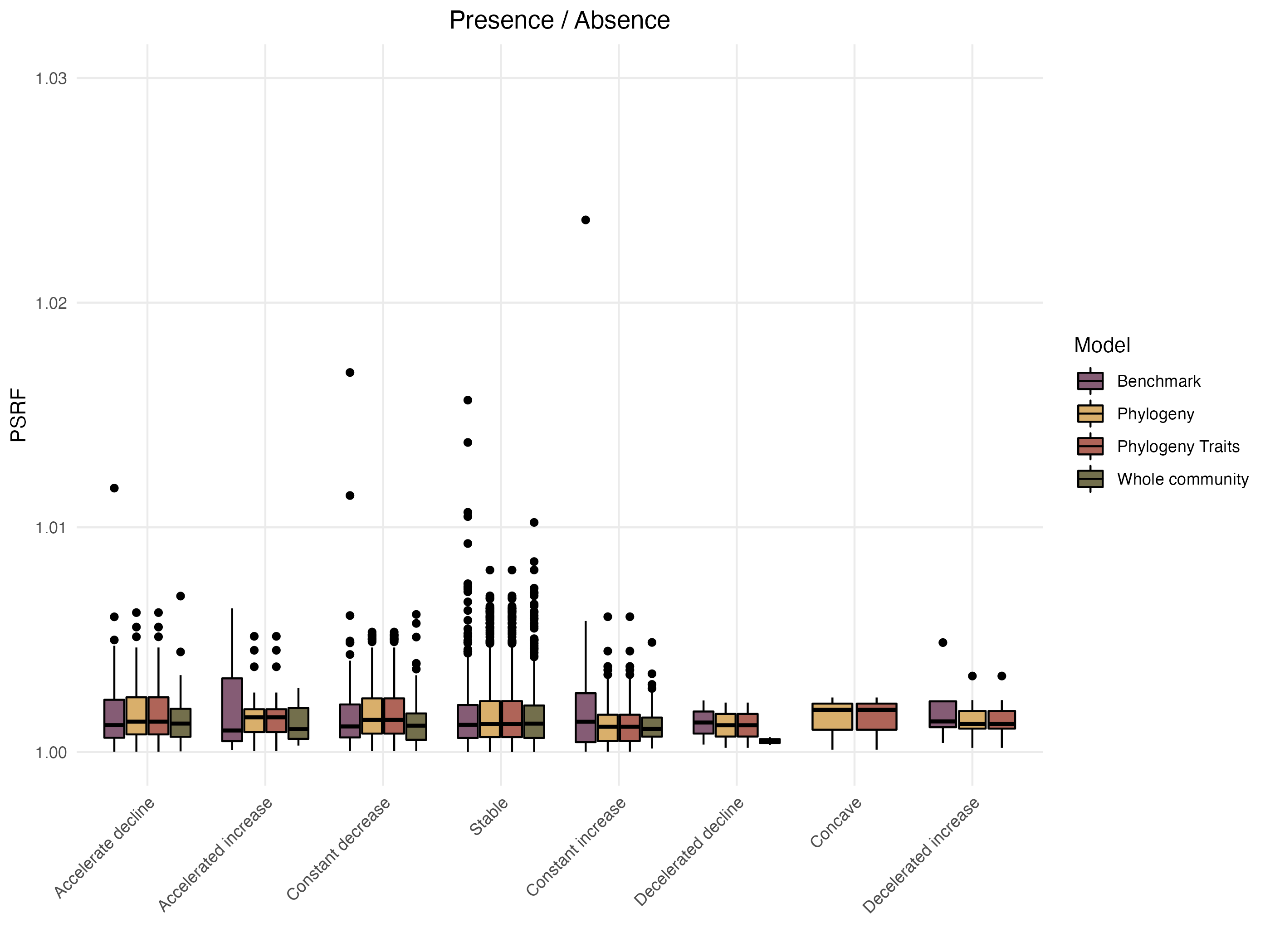

Figure 16: Distribution of the Potential scale reduction factors (PSRF) as a function of the different shapes of the response curves classified following the methodology proposed by Rigal *et al.* (2020) (see section “Assessing model performance and interpretability” section for more details on the calculation methodology). Results for models fitted with presence/absence data.

### Appendix C - Complementary Results

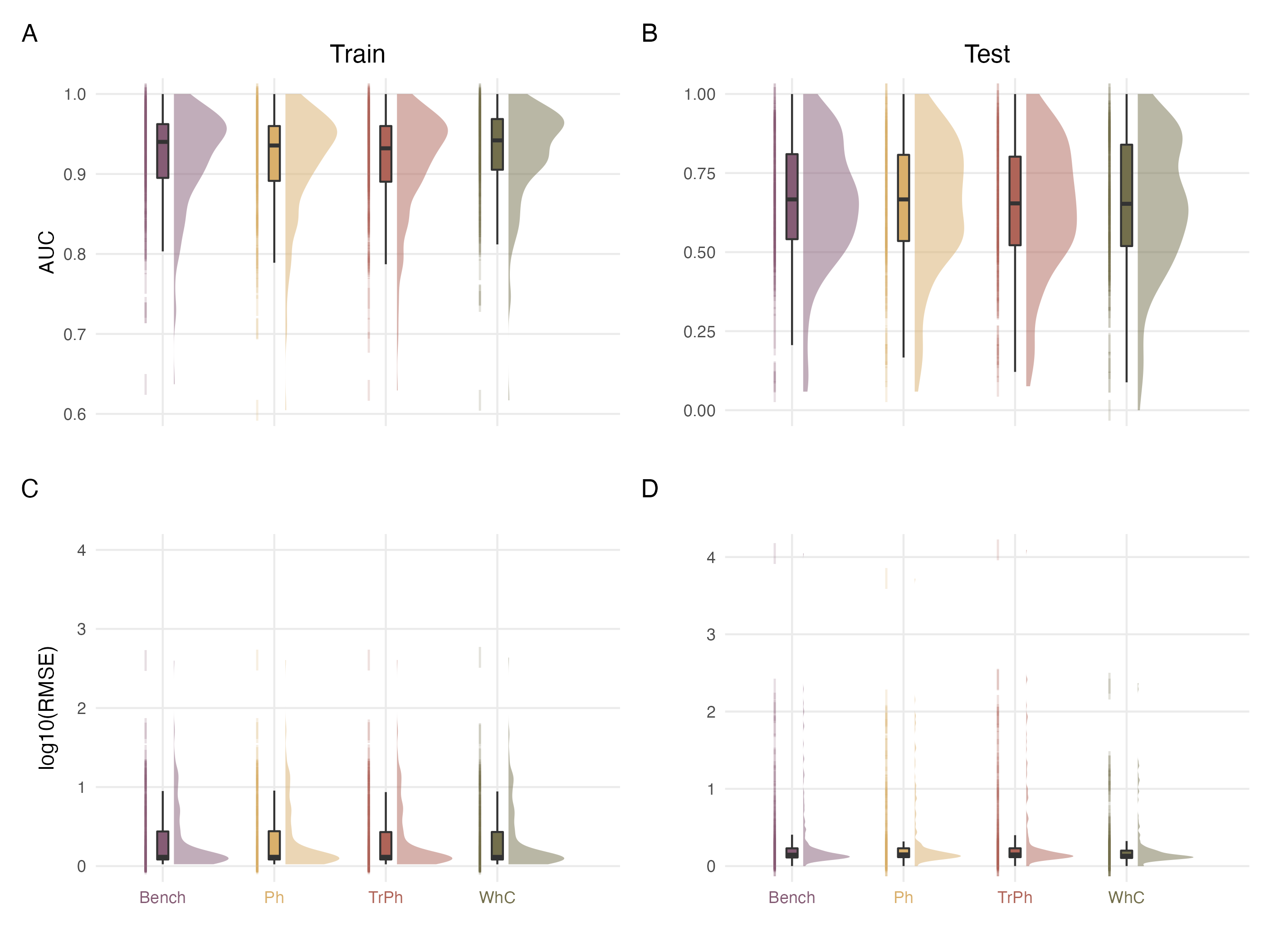

Figure 17: Comparison of explanatory (left column; Train set) and predictive (right column; Test set) performance of the different model structures fitted on presence/absence (top panels) or abundance (bottom panels) data.

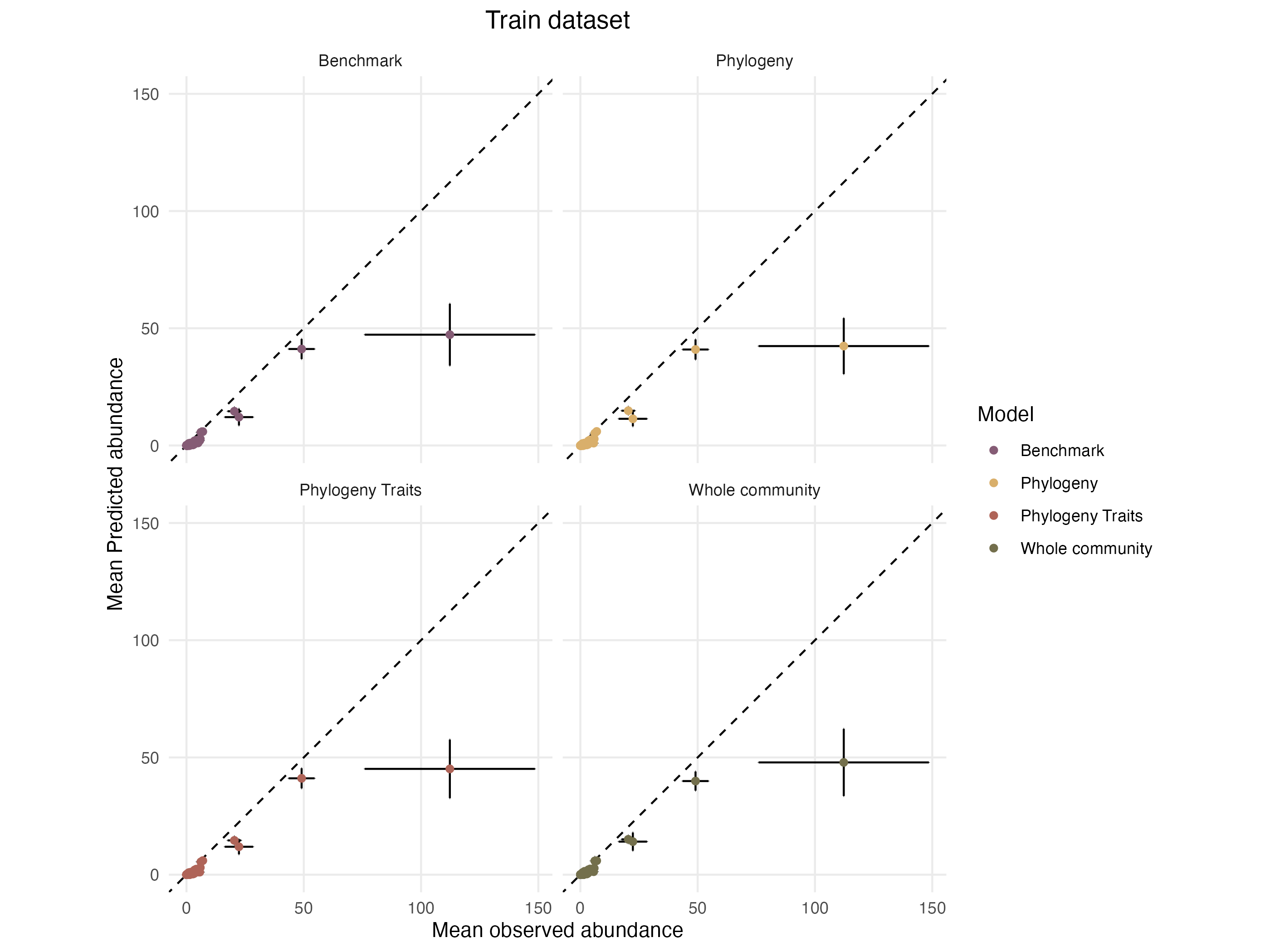

Figure 18: Mean predicted abundance as a function of mean observed abundance in the training dataset. Each species is represented by a dot, the error bars on each point indicate the standard error around the mean relative to each axis.

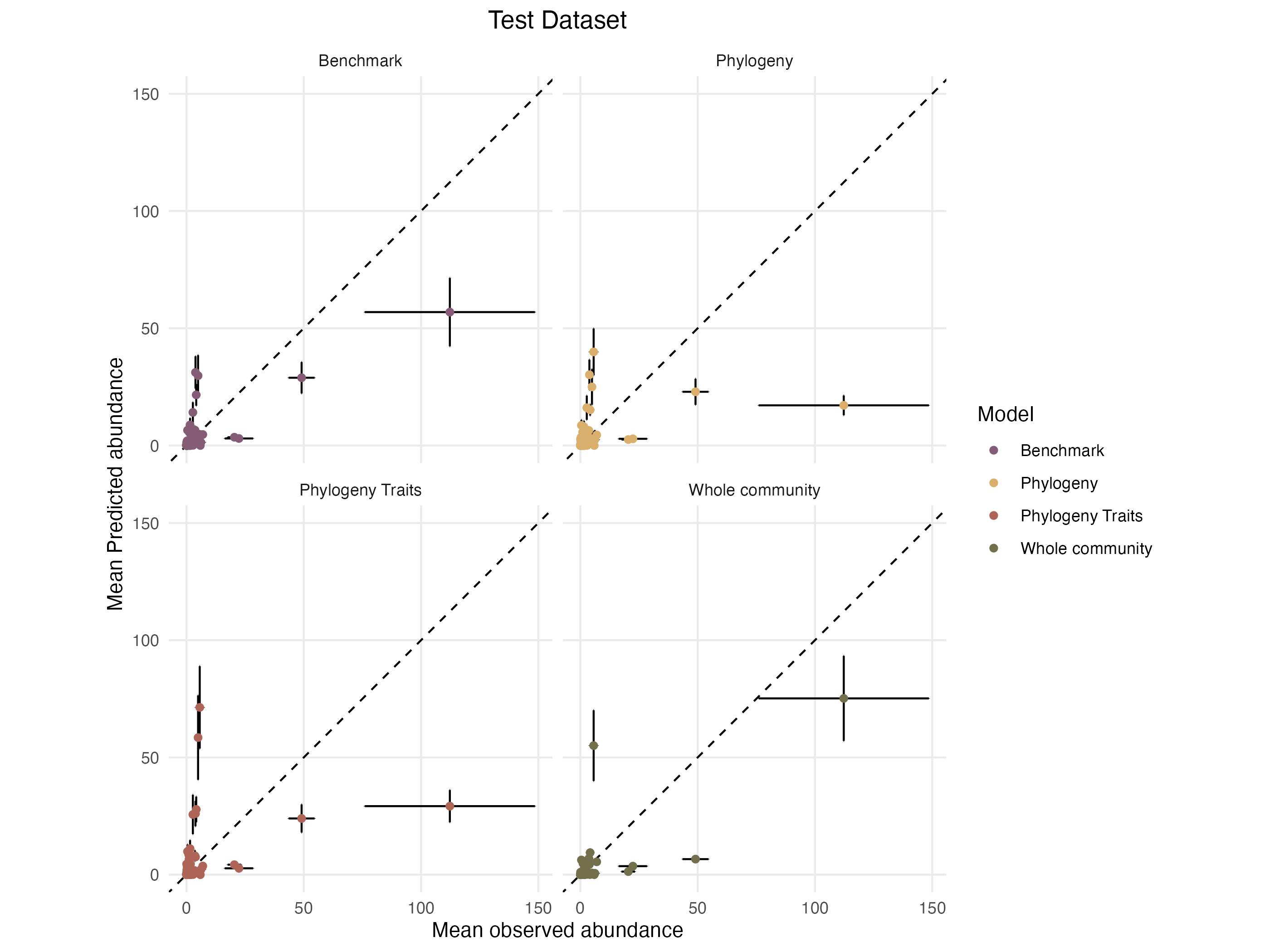

Figure 19: Mean predicted abundances as a function of mean observed abundances in the training dataset. Each species is represented by a dot, the error bars on each point indicate the standard error around the mean relative to each axis.

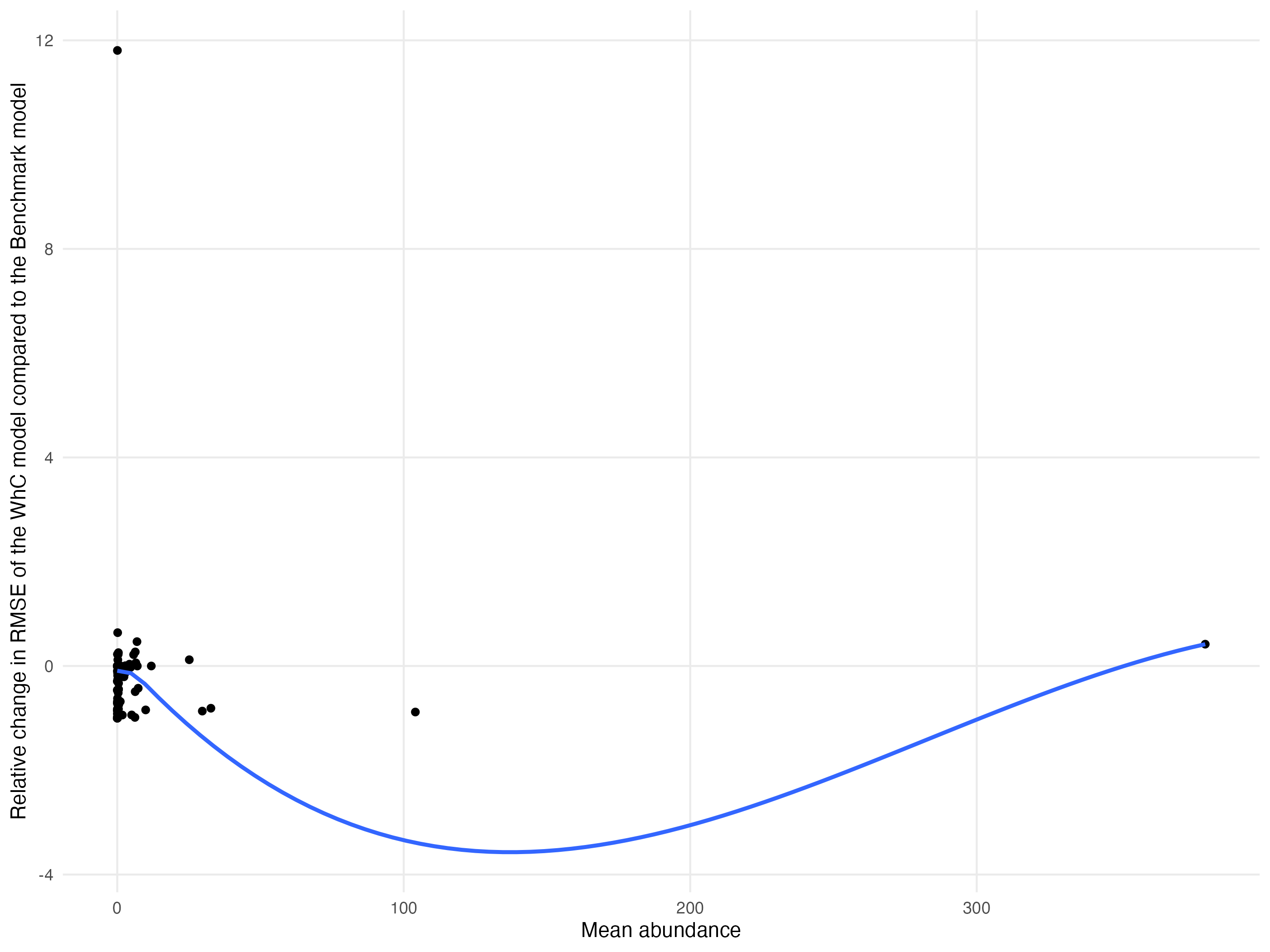

Figure 20: Relationship between the relative improvement in RMSE for the WhC model compared to the Bench model and the mean abundance of species in the training dataset. Each dot represents a species. The blue line represents a fit obtained from a LOESS regression.

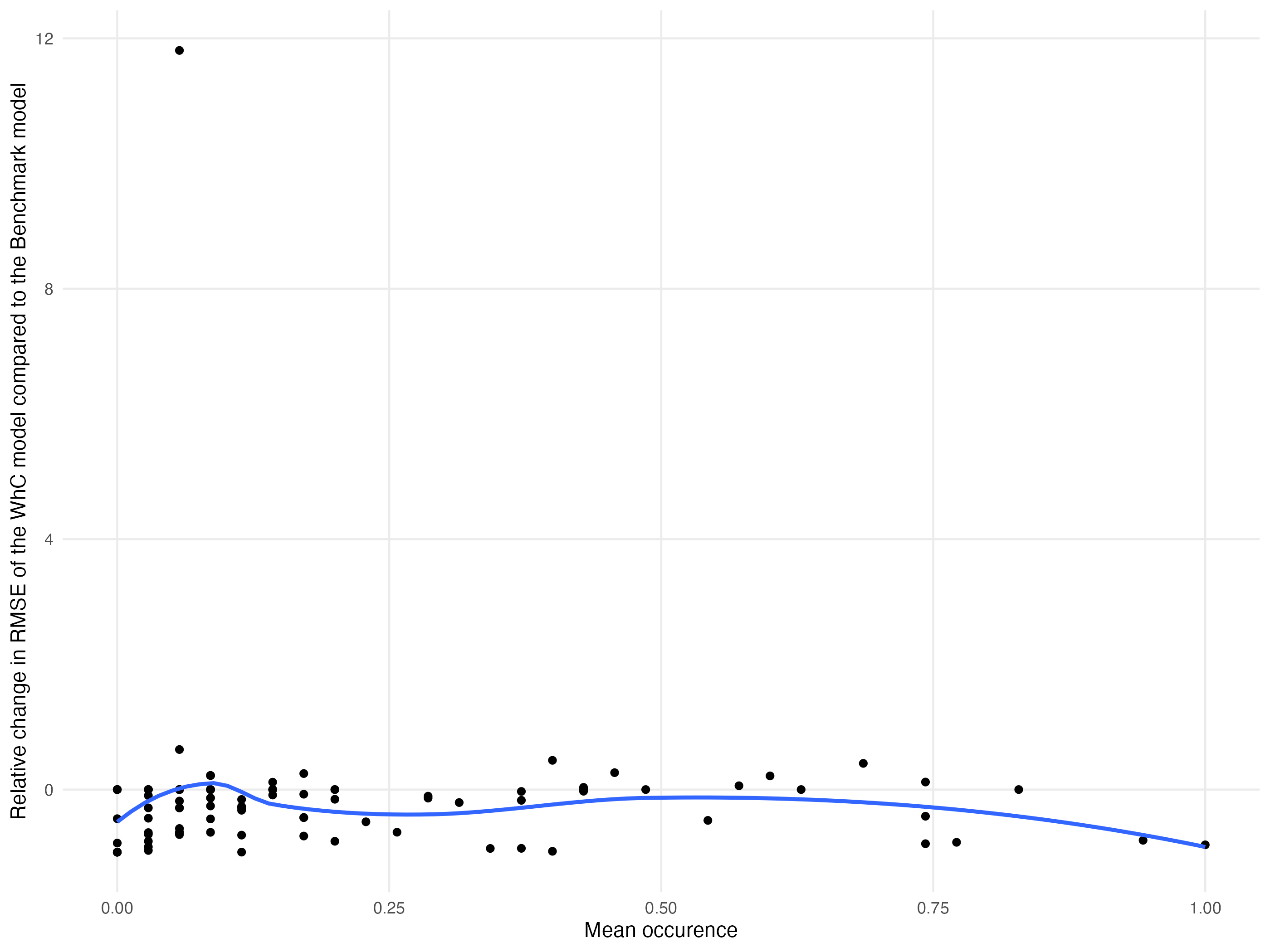

Figure 21: Relationship between the relative improvement in RMSE for the WhC model compared to the Bench model and the mean occurrence of species in the training dataset. Each dot represents a species. The blue line represents a fit obtained from a LOESS regression.

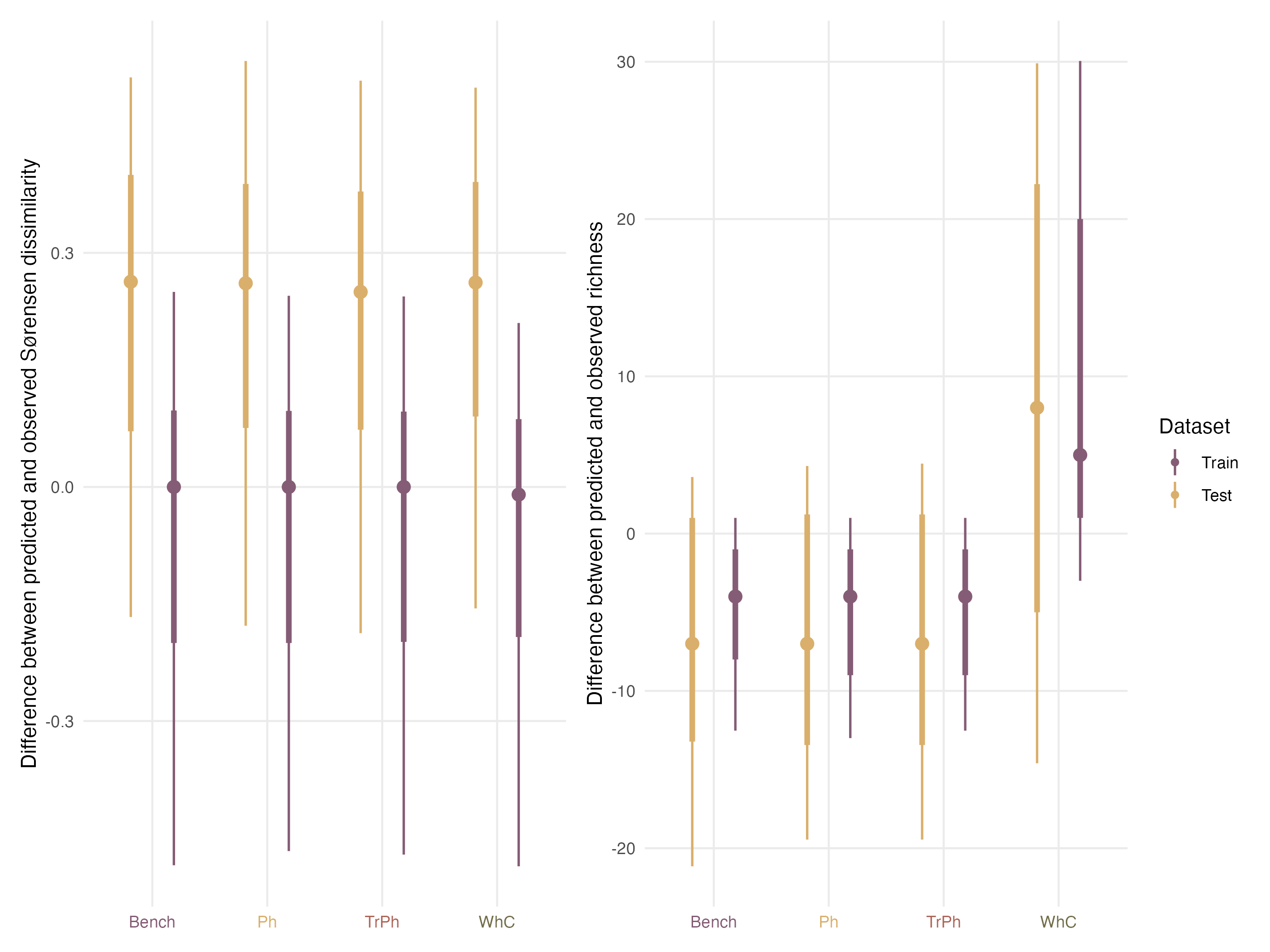

Figure 22: Comparison of model performances with regards to their ability to predict community structures when fitted with presence/absence data for the train (purple) and test (yellow) dataset. The left column indicates for each model the difference between the pairwise dissimilarities computed on the observed assemblages and those computed on the predicted community. The right column presents differences in species richness between the observed and predicted assemblages.

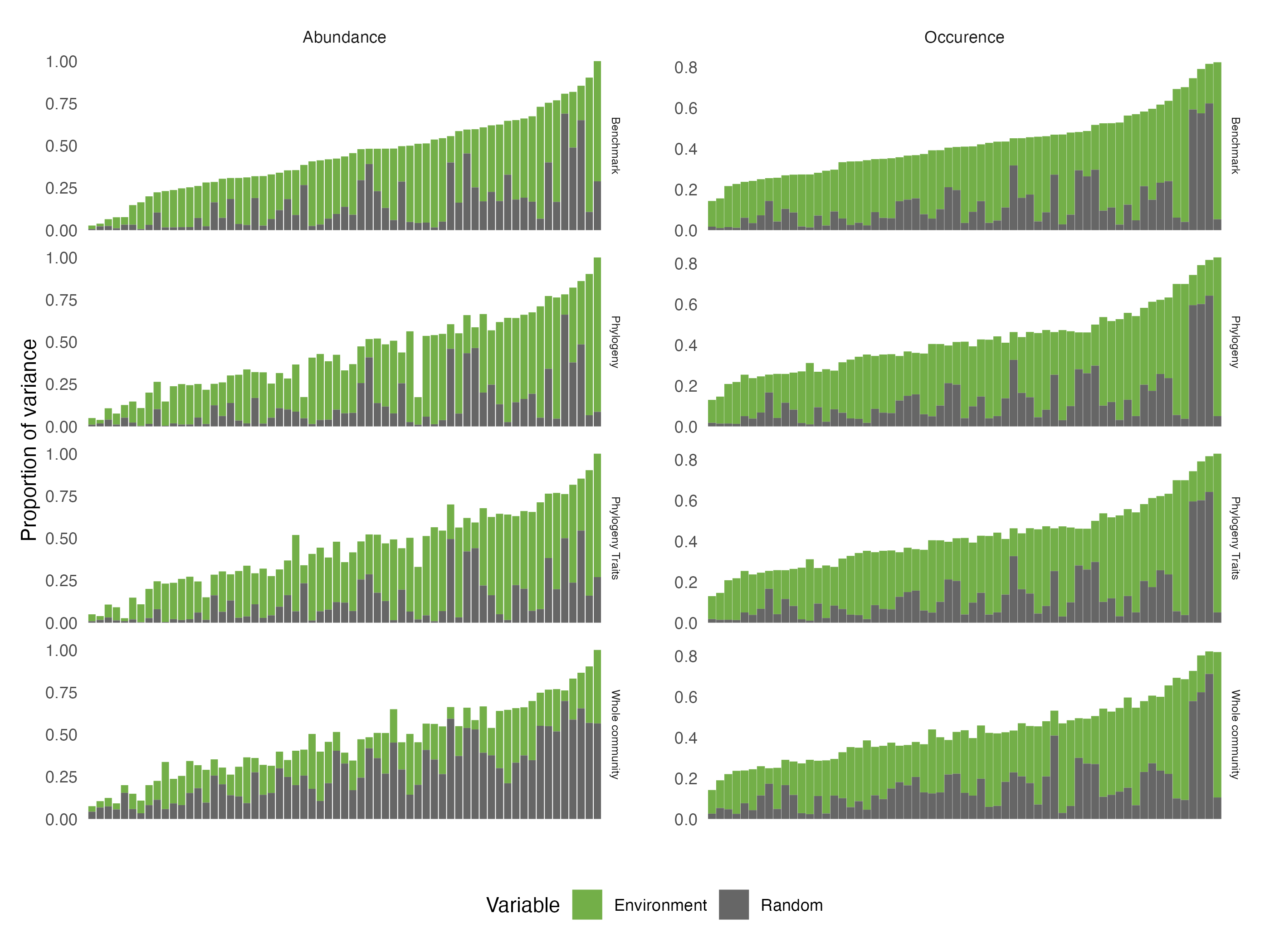

Figure 23: Comparison across the four alternative model structures of the total amount of variance explained by (1) the environmental variables (Environment) and (2) the three random effects (Random), for each species (x-axis). Results are presented for the models fitted with abundance (left) and presence/absence (right) data. Species are ordered by increasing order of total variance explained by the benchmark model.

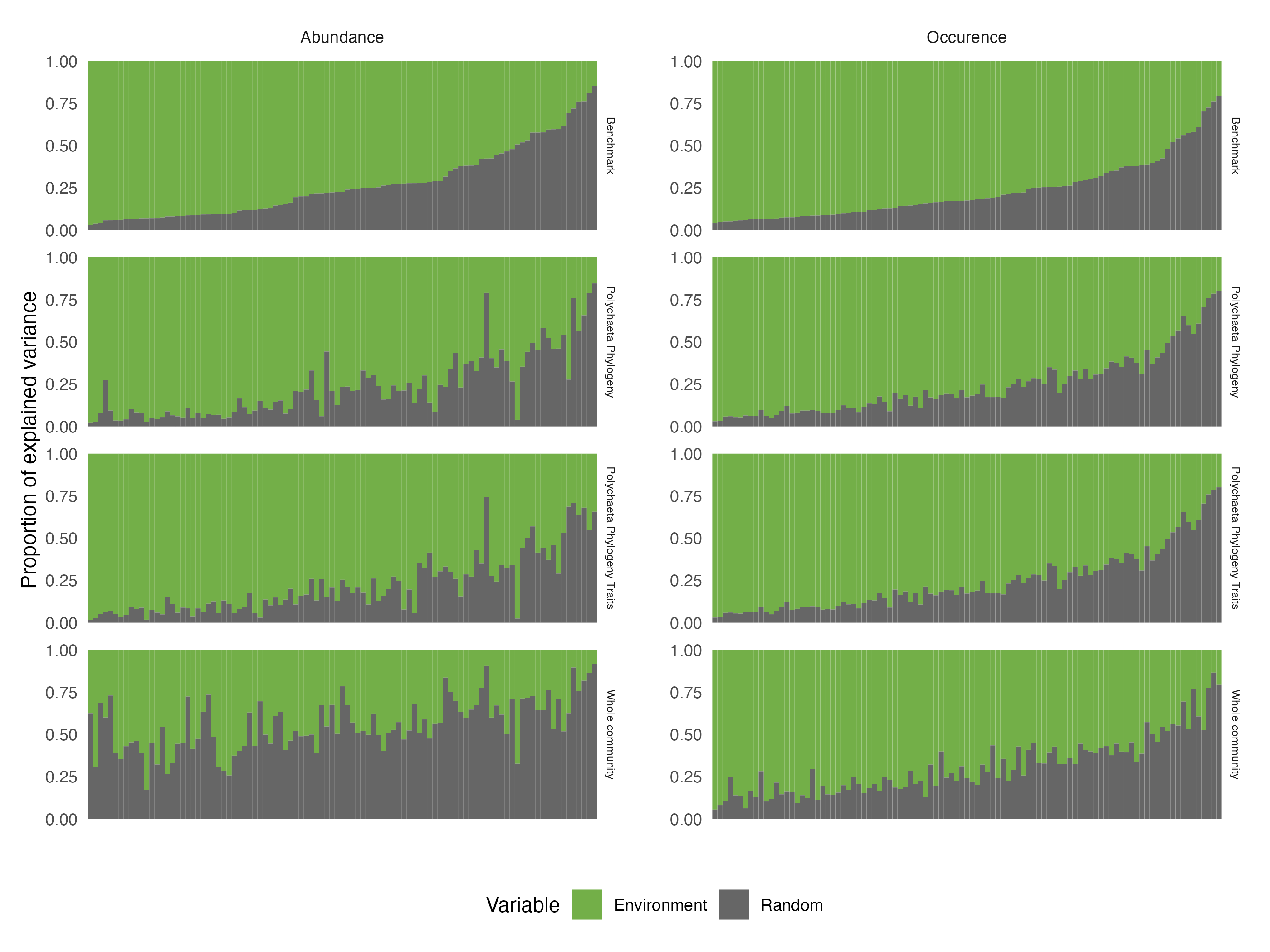

Figure 24: Comparison across the four alternative model structures of the fraction of variance explained by (1) the environmental variables (Environment) and (2) the three random effects (Random) for each species (x-axis) . Results are presented for the models fitted with abundance (left) and presence/absence (right) data. Species are ordered by decreasing order of variance explained by the environment for the benchmark model.

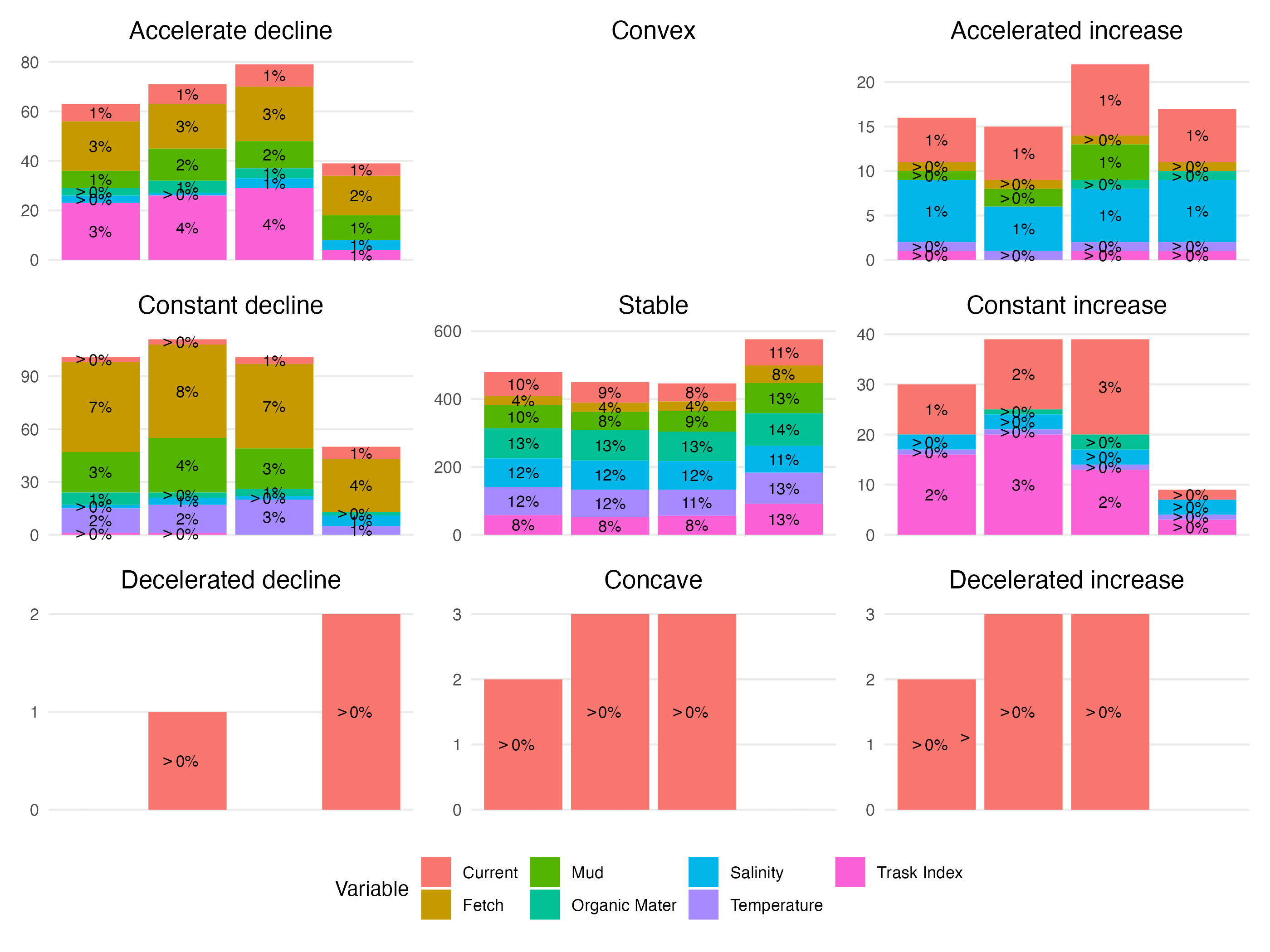

Figure 25: Same figure as Fig. 4 in the main text for models fitted to abundance data. Number (y-axis) and proportion (indicated above individual bars, rounded to the nearest integer) of response curves (i.e. one for each species-predictor combination) according to the nomenclature (nine shapes highlighted by the black curve in each panel) proposed by Rigal *et al.* (2020). Results are presented for different model structures: from left to right the Benchmark (Bench), the phylogeny (Ph), the traits & phylogeny (TrPh), and the whole community (WhC) models. Each bar is coloured by the relative contribution of each environmental covariate to this particular shape. For illustrative purposes, note that the scale of variation on the y-axis differs across panels.

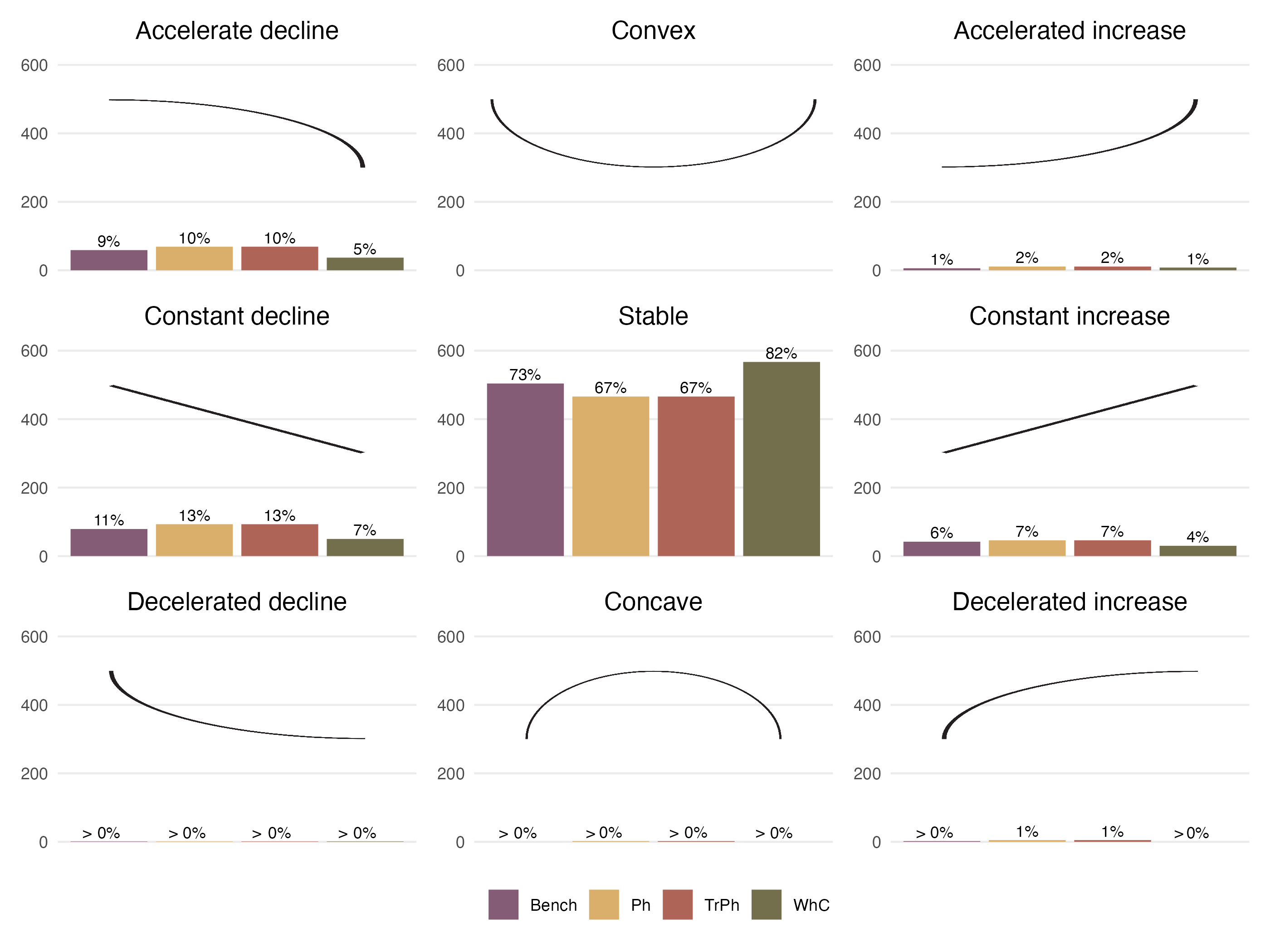

Figure 26: Same figure as Fig. 4 in the main text but for models fitted to presence/absence data. Number (y-axis) and proportion (indicated above individual bars, rounded to the nearest integer) of response curves (i.e. one for each species-predictor combination) according to the nomenclature (nine shapes highlighted by the black curve in each panel) proposed by Rigal *et al.* (2020). Results are presented for the different structures: purple for the Benchmark (Bench), yellow for phylogeny (Ph), red for traits & phylogeny (TrPh), and green for the whole community (WhC) model.

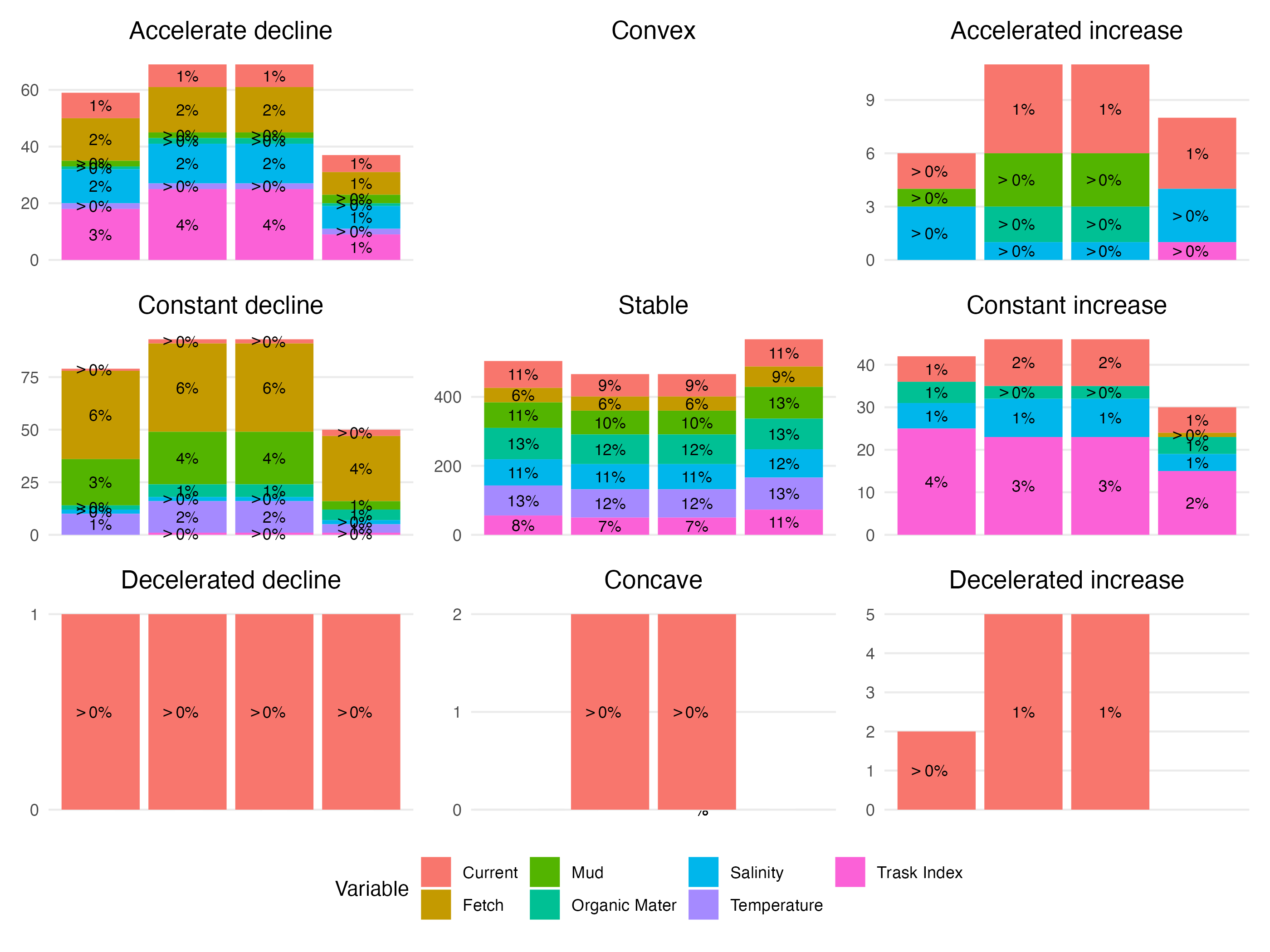

Figure 27: Same figure as Fig. 25 but for models fitted on presence/absence data. Number (y-axis) and proportion (indicated above individual bars, rounded to the nearest integer) of response curves (i.e. one for each species-predictor combination) according to the nomenclature (nine shapes highlighted by the black curve in each panel) proposed by Rigal *et al.* (2020) for different presence/absence model structures. Each bar is coloured by the relative contribution of each environmental covariate to this particular shape. For illustrative purposes, note that the scale of variation on the y-axis differs across panels.

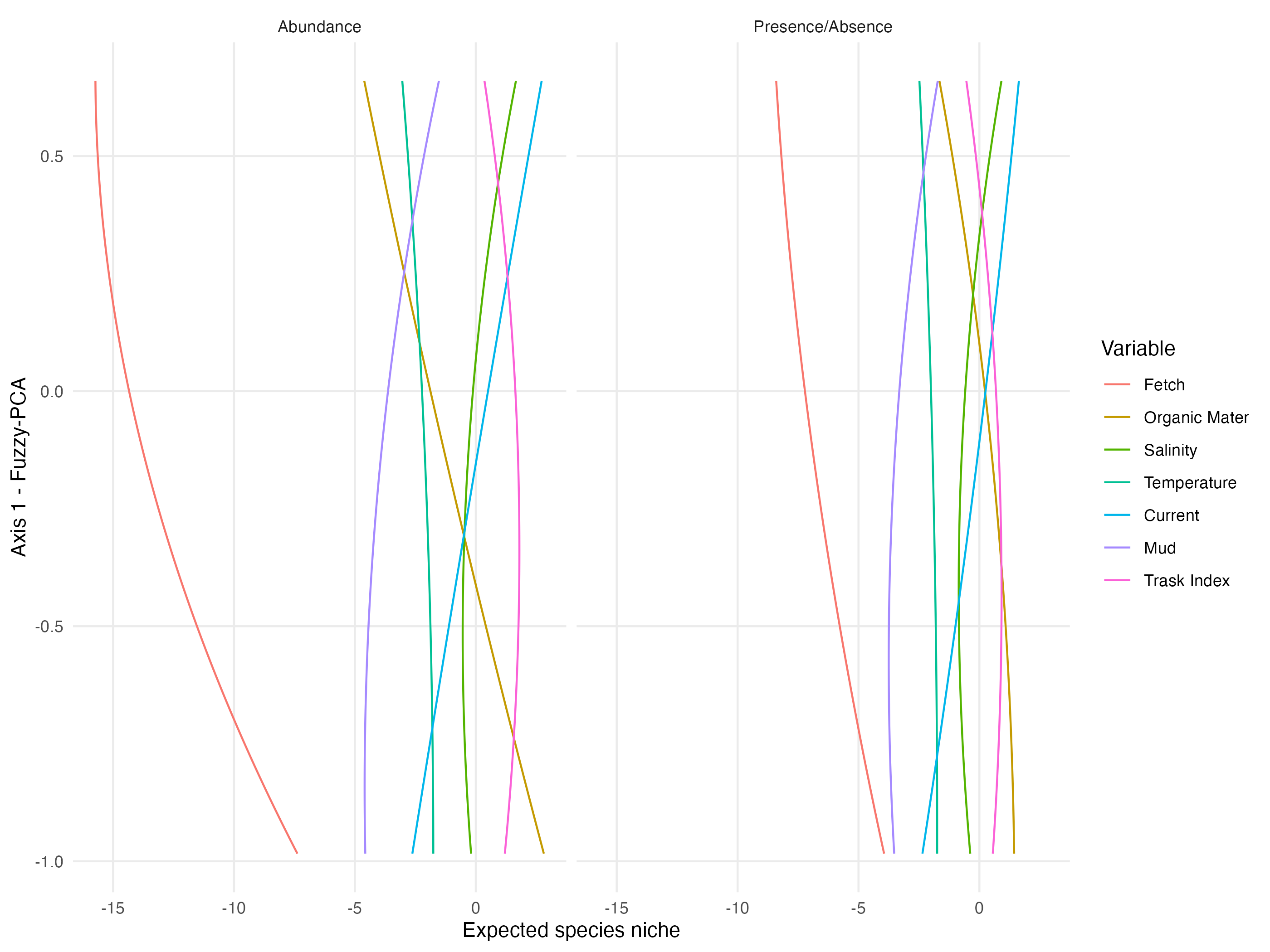

Figure 28: Relationships between species’ position along the first axis of the fuzzy PCA (sessile microphagous-mobile macrophagous gradient) and the different environmental variables used in the models (fitted with abundance data in the left panel, and with presence/absence data in the right panel). Relationships are derived from the regression coefficients estimated for the TrPh model (γ coefficients in HMSC; Ovaskainen & Abrego (2020)).The lines are fitted quadratic regressions representing the average response across the different species. As an example of interpretation, the red lines in both graphs indicate that sessile microphagous species are more negatively influenced (lower abundance, low probability for presence) by fetch than macrophagous mobile species.

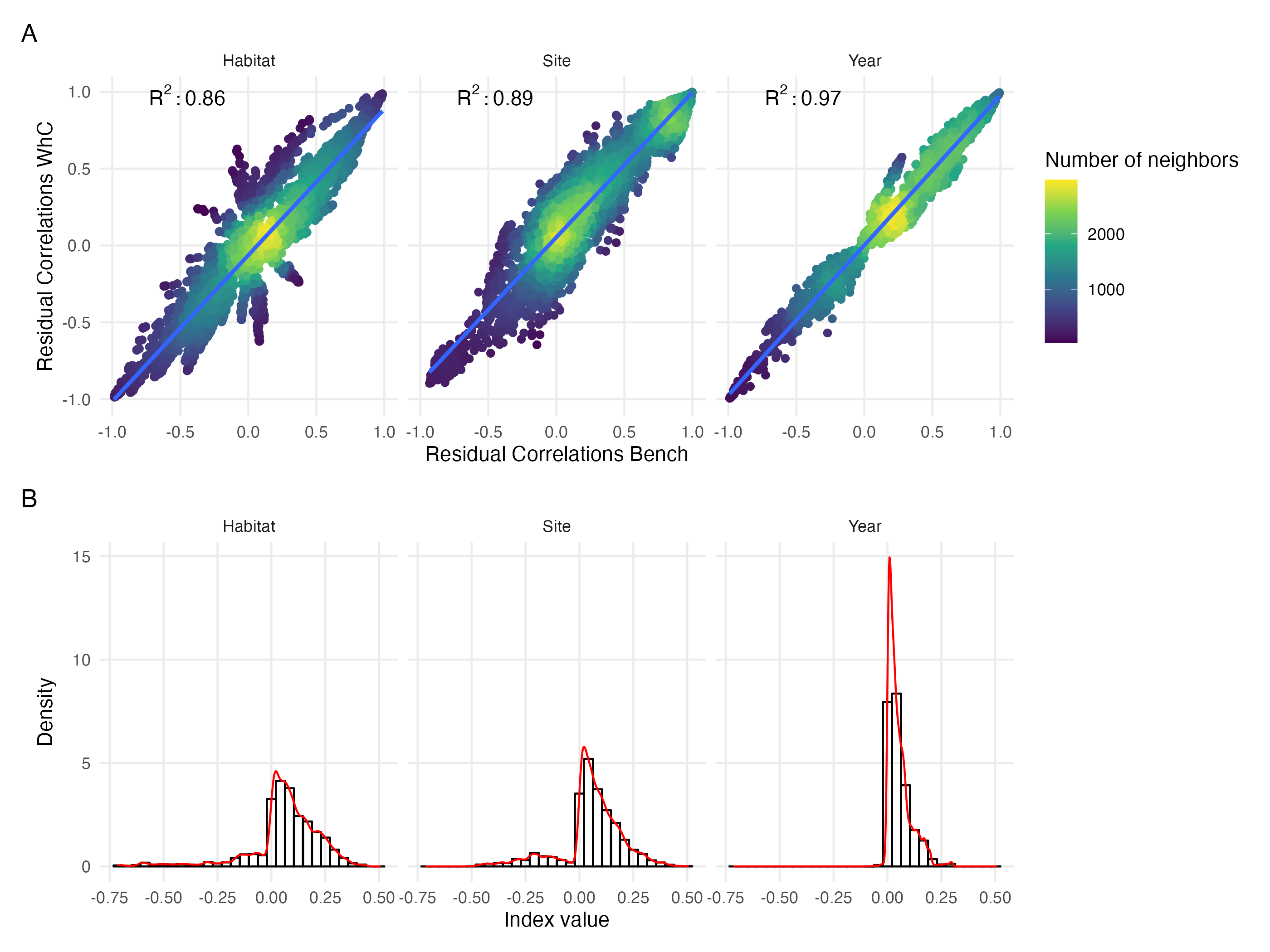

Figure 29: Same figure as Fig. 5 in the main text but for models fitted on presence/absence data. (A) Comparison of residual correlations associated with the three random effects estimated by the Whole Community Model (y-axis) and the Benchmark model (x-axis). The colour scale highlights the density of points in each scatter plot. (B) Distribution of the index measuring change in sign (sign change left to the zero line, no change to the right) and magnitude (higher departure from the zero line indicates higher difference) between residual correlations estimated by the whole community model and the benchmark model for the three random effects (Habitat, Site, Year).
